# Visualization of the HIV-1 Env Glycan Shield Across Scales

**DOI:** 10.1101/839217

**Authors:** Zachary T. Berndsen, Srirupa Chakraborty, Xiaoning Wang, Christopher A. Cottrell, Jonathan L. Torres, Jolene K. Diedrich, Cesar A. López, John R. Yates, Marit J. van Gils, James C. Paulson, S Gnanakaran, Andrew B. Ward

## Abstract

The dense array of N-linked glycans on the HIV-1 Envelope Glycoprotein (Env), known as the “glycan shield”, is a key determinant of immunogenicity, yet intrinsic heterogeneity confounds typical structure-function analysis. Here we present an integrated approach of single-particle electron cryomicroscopy (cryo-EM), computational modeling, and site-specific mass-spectrometry (MS) to probe glycan shield structure and behavior at multiple levels. We found that dynamics lead to an extensive network of inter-glycan interactions that drive the formation of higher-order structure within the glycan shield. This structure defines diffuse boundaries between buried and exposed protein surface and creates a mapping of potentially immunogenic sites on Env. Analysis of Env expressed in different cell lines revealed how cryo-EM can detect subtle changes in glycan occupancy, composition, and dynamics that impact glycan shield structure and epitope accessibility. Importantly, this identified unforeseen changes in the glycan shield of Env obtained from expression in the same CHO cell line used for GMP production. Finally, by capturing the enzymatic deglycosylation of Env in a time-resolved manner we found that highly connected glycan clusters are resistant to digestion and help stabilize the pre-fusion trimer, suggesting the glycan shield may function beyond immune evasion.

**Significance Statement:** The HIV-1 Env “glycan shield” masks the surface of the protein from immune recognition, yet intrinsic heterogeneity defies a typical structure-function description. Using a complementary approach of cryo-EM, computational modeling, and mass-spectrometry we show how heterogeneity and dynamics affect glycan shield structure across scales. Our combined approach facilitated the development of new cryo-EM data analysis methods and allowed for validation of models against experiment. Comparison of Env across a range of glycosylation states revealed how subtle differences in composition impact glycan shield structure and affect the accessibility of epitopes on the surface. Finally, time-resolved cryo-EM experiments uncovered how highly connected glycan clusters help stabilize the pre-fusion trimer, suggesting the glycan shield may function beyond immune evasion.

## Introduction

The Human Immunodeficiency Virus Type 1 (HIV-1) Envelope Glycoprotein (Env) is the sole antigen on the surface of the virion and has evolved several tactics for evading the adaptive immune system, chief among which is extensive surface glycosylation (1–3). Env has one of the highest densities of N-linked glycosylation sites known, with glycans accounting for ~1/2 the mass of the molecule (4–8). This sugar coat, or “glycan shield,” is common among viral fusion proteins and is believed to be a primary hurdle in the development of neutralizing antibodies against Env during infection and vaccination (9, 10). Therefore, arriving at a consistent and general description of its structure and dynamics may prove necessary in designing an effective Env-based immunogen and be of broader importance towards understanding the structure and function of other densely glycosylated proteins.

N-linked glycans can be highly dynamic, however, and can vary substantially in length, connectivity, and chemical composition (11, 12). On the Env ectodomain surface there are ~95 glycosylation sites on average (6), and mass-spectrometry analysis (MS) has revealed that compositional heterogeneity exists between sites as well as at the same site between different copies of Env (13). While heterogeneity and dynamics are critical for immune evasion, they are incompatible with a typical structure-function description. When the first structures of the fully glycosylated Env were solved with single-particle electron cryomicroscopy (cryo-EM) (14, 15) most glycans were not resolved beyond the first or second sugar ring except when stabilized by antibodies. This suggested that the glycan shield is highly dynamic, however, due to the missing high-resolution details, structural analysis remained focused on Env-antibody interactions, while experimental investigations into the glycan shield shifted primarily towards chemical analysis via MS (11). One crystallography study specifically addressing glycan shield structure was recently published and the data shows evidence for highly stabilized glycans engaging in a wide range of inter-glycan contacts (6), which is at odds with the cryo-EM results. Such a disagreement between the two techniques highlights the difficulty in studying the glycan shield and the need for more focused experiments.

Computational modeling, in particular molecular dynamics (MD) simulations, has proven useful as a complementary approach for characterizing glycoprotein structure and dynamics (16–19), including Env (6, 20–22). Three sets of atomistic dynamics simulations of fully glycosylated Env trimers were recently reported and these predicted that dynamics would lead to extensive shielding of antigenic surface and the formation of complicated networks among interacting glycans (6, 21, 22). In one study, glycans were observed clustering into distinct micro-domains over long timescales, and it was found that neutralizing antibodies preferentially target the interfaces between these domains, suggesting the glycan shield may possess biologically relevant large-scale structure. Although MD is a widely accepted method for modeling glycoproteins, atomistic simulations for systems as large as Env require substantial computational resources to access biologically relevant spatial and temporal scales and can suffer from initial model bias. Therefore, more computationally efficient methods for modeling fully glycosylated Env are needed. Furthermore, there are still no established methods for experimentally validating the results of these simulations.

Here, we describe an integrated approach combining cryo-EM, computational modeling, and site-specific MS aimed at illuminating glycan shield structure and behavior at multiple levels. First, we introduce a model system for our cryo-EM experiments and describe the complete structure of the native Env glycan shield for the first time using scale-space and 3-D variability analysis. In parallel, we developed a high-throughput modeling pipeline for rapidly generating diverse ensembles of fully glycosylated Env at atomistic resolution, enabling quantification of glycan-specific geometric properties and concerted behavior within the glycan shield as a whole. We then used these ensembles as ground-truth for the creation of simulated cryo-EM data, which facilitated the development of new analysis tools and enabled the validation of theoretical models against cryo-EM experiments.

With our integrated approach, we show that the glycan shield is highly dynamic but exhibits variation in dynamics between glycans due in part to crowding and other geometric and energetic constraints. We found that dynamics give rise to a network of inter-glycan interactions that drive the formation of higher-order structure within the glycan shield. This blurry low-resolution structure creates diffuse boundaries between buried and exposed protein surface that define potential sites of vulnerability. Using Env expressed in three common cell lines, we show how differences in glycan composition and occupancy can be detected by cryo-EM and result in changes to glycan shield structure and dynamics that affect the accessibility of epitopes on the surface. To better inform our results we present site-specific mass-spectrometry data for the same samples and demonstrate that cryo-EM can complement such studies. Finally, by exposing Env to endoglycosidase digestion and capturing reaction intermediates with cryo-EM in a time-resolved manner, we found that highly connected glycan clusters are resistant to digestion and act to stabilize the pre-fusion trimer structure, implying the glycan shield may function beyond immune evasion.

## Results

### Env in complex with a base-specific antibody as a model system for cryo-EM analysis of the native glycan shield

We choose BG505 SOSIP.664 as a model system to demonstrate our approach because it is the first and most widely studied of the soluble pre-fusion stabilized Envs (23, 24). Native Env is a transmembrane protein that forms a trimer of dimers composed of two gene products, gp120 and gp41, which are responsible for receptor binding and membrane fusion respectively [add citation]. The acronym *SOSIP* is in reference to the engineered disulfide bonds between gp120 and gp41 and the isoleucine to proline mutation in the HR1 helix, which act to stabilize the pre-fusion conformation, while 664 is the final residue in truncated ectodomain (23). BG505 SOSIP.664 is also currently in the first ever human clinical trials of a trimeric subunit-based HIV vaccine (https://clinicaltrials.gov/ct2/show/NCT03699241). In Fig. 1 we present the cryo-EM reconstruction and refined atomic model of BG505 SOSIP.664 purified from HEK293F cells in complex with three copies of the fragment antigen binding domain (Fab) of the non-neutralizing antibody, RM20A3. This complex will be referred to as BG505_293F from here on. The RM20A3 Fab was included because it improves orientation bias, as illustrated by the complete and near-uniform angular distribution (Fig. 1*C*), and it binds to a non-native epitope at the base of the SOSIP trimer that is devoid of glycans (Fig. 1*D*). This leaves the native glycan shield undisturbed and means that RM2OA3 Fab binding is independent of glycosylation, making the SOSIP/base-Fab complex ideal for this study.

**Fig 1.**
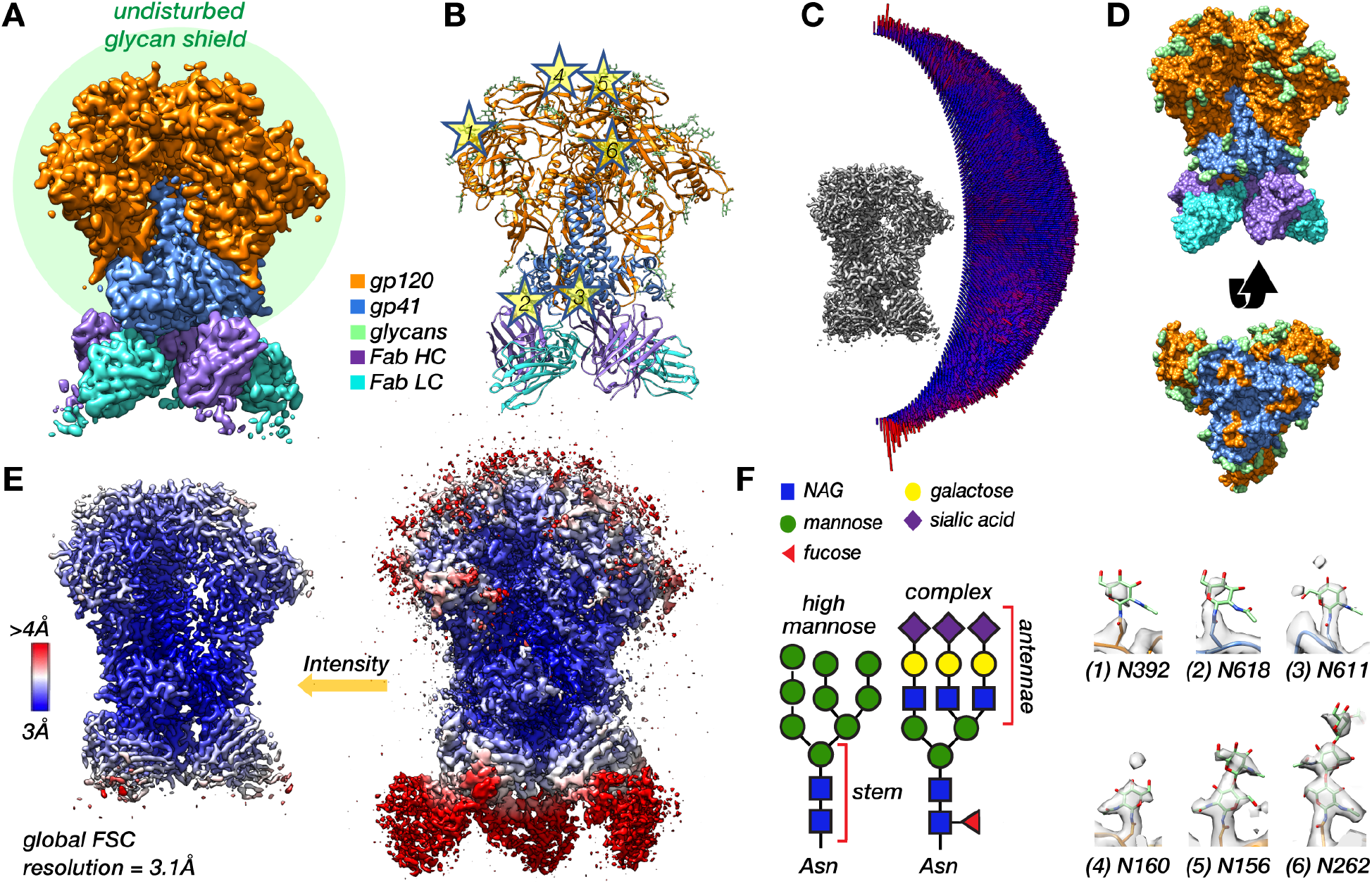
Soluble SOSIP in complex with a base-specific Fab as a model system for cryo-EM analysis of the native HIV-1 Env glycan shield. **(A)** Segmented cryo-EM map of BG505 SOSIP.664 in complex with three copies of the RM20A3 Fab. The green circle is meant to represents the glycan shield. **(B)** Refined atomic model with stars highlighting several N-linked glycans. **(C)** 3-D angular distribution histogram. **(D)** Surface representation of the refined atomic model viewed from the side and from the bottom with the Fab chains removed. (**E**) Sharpened 3.1Å-resolution cryo-EM map at high and low threshold colored by local resolution. **(F)** SNFG depiction (74) of a representative high-mannose and complex type glycan (L). Cryo-EM map density for 6 representative glycans from the refined atomic model (numbers corresponding to the stars in panel B).

The global resolution of this C3-symmetric reconstruction determined by Fourier shell correlation (FSC) is ~3.1Å (*SI Appendix*, Fig. S1*A*), and the bulk of the protein is at or near this resolution (Fig. 1*E*). ~80% of Env residues could be confidently built into this map, while most glycans are ill-defined beyond the core N-acetylglucosamine (NAG), accounting for only ~15% of the total predicted glycan mass (Fig. 1). Glycans located on the poorly resolved flexible loops (N185e, N185h, N398, N406, N411, N462) could not be identified at all, however, site-specific MS data confirmed they are indeed glycosylated (25) (*SI Appendix*, Fig. S2*A*). N-linked glycans on Env can be roughly classified into two types, high-mannose and complex, based on the extent of intracellular processing, and they differ in composition and structure (one possible configuration of each type is depicted in Fig. S 1F). Even if the glycans were better resolved, averaging of individual projections during 3-D reconstruction means we cannot uniquely identify the specific type of glycan at most sites because they possess a mixture of several different types (26) (*SI Appendix*, Fig. S2*A*). Of those we could identify, a few were found to be near-uniformly processed (designated as >75% complex), for example the glycans at N611 and N618 on gp41, and it should be possible to confirm this by the presence of a core fucose residue. However, we were unable to identify clear density for fucose at any of these sites (Fig. 1*F*). Together, these observations show that the glycan shield is highly dynamic with respect to the protein core but exhibits some differences in ordering between glycans, and that it is not possible to distinguish between the different types of glycans from a single cryo-EM map when the glycans are not stabilized by antibodies.

### Scale-space and 3-D variance analysis reveal interconnectivity and higher-order structure within the glycan shield

Although we lack the resolution to build precise atomic models, there is still a great deal of information about glycan shield structure and dynamics that can be gleaned from the cryo-EM data. At low isosurface thresholds noise appears surrounding Env’s well resolved protein core where the missing glycan mass should be (Fig. 1*E*). To look for meaningful structure within this noisy signal we analyzed properties of the map across a range of scales by progressively smoothing it with a Gaussian filter of increasing standard deviation (SD). Resolution in the glycan shield falls off with distance from the protein surface, therefore as the map is smoothed more and more glycan signal should become visible until the map encompasses the majority of the space occupied by the ensemble of configurations. This process could be captured by measuring the volume of the map across the scale-space. To calculate the volume, however, one must first choose an appropriate threshold. For our analysis we implemented an auto-thresholding method based a simple measure of topological connectivity described in the *SI Appendix* (*SI Appendix*, Fig. S3*A-F*). We refer to the resulting intensity as the *noise threshold* since it represents the lowest threshold before the appearance of noise. Fig. 2*A* shows the noise threshold and volume of the map at the noise threshold as a function of Gaussian filter width. Both curves have a sigmoidal shape; once the map is sufficiently smoothed it undergoes a period of rapid expansion in volume and reduction in noise threshold that is presumably capturing the appearance of signal from the poorly resolved regions of the map. The curves then plateau around ~1.5-2 SD, suggesting there is negligible gain in signal with further smoothing.

**Fig 2.**
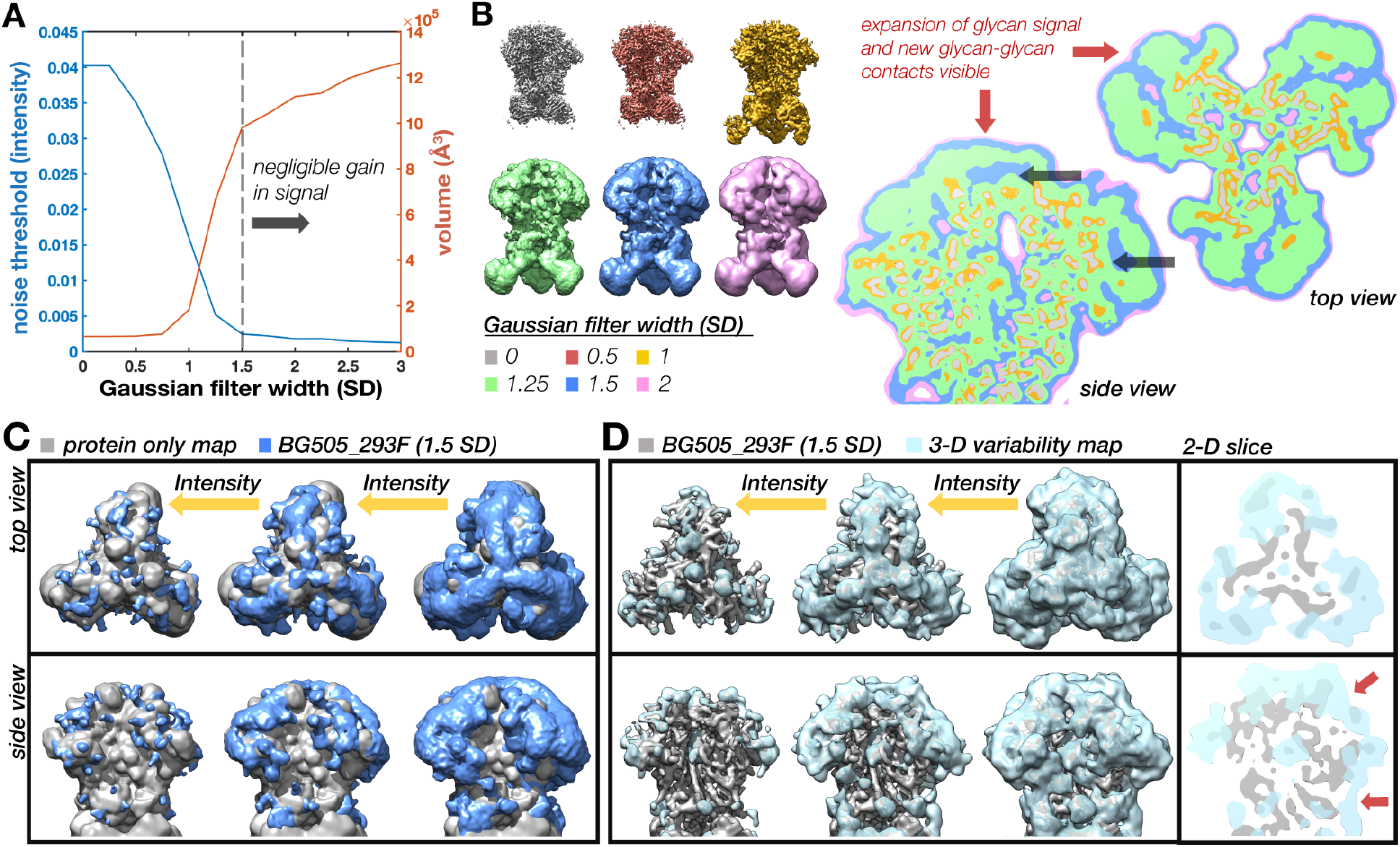
Scale-space and 3-D variability analysis reveal low resolution structure in the glycan shield. **(A)** Noise threshold and volume at the noise threshold as a function of Gaussian filter width measured in standard deviation (SD) showing the emergence of glycan signal at low resolutions and plateau around 1.5 SD. **(B)** BG505_293F cryo-EM map at 6 representative scales along with 2-D slices through the maps overlaid on top of one another showing the emergence of glycan signal and new glycan-glycan contacts visible at low resolutions and thresholds (red arrows). Black arrows are pointing to protein surfaces occluded by glycans. **(C)** Gaussian filtered (1.5 SD) BG505_293F map (blue) visualized at three intensity thresholds along with a protein-only map (gray). **(D)** SPARX 3-D variability map (light blue) visualized at three intensity thresholds along with the BG505_293F map (gray) and 2-D slices through the top and side of the map with red arrows highlighting glycan-glycan contacts.

Examination of these filtered maps confirms the progressive emergence of more structure in the glycan shield (Fig. 2*B*). By taking 2-D slices through six representative maps and displaying them on top of one another we see how glycosylated surfaces undergo substantially more dilation (red arrows) than the non-glycosylated surfaces. Once the isosurface has expanded to include signal from glycan antennae, neighboring glycans begin to merge, creating a canopy-like effect that occludes underlying protein surfaces (black arrows). This implies that neighboring glycans sample overlapping volumes and have the potential to interact. Also apparent is the negligible gain in signal moving from 1.5 to 2 SD (blue and pink). We therefore concluded that the 1.5 SD Gaussian filtered map is the ideal scale for interpreting glycan shield structure in this reconstruction. Visualization of this filtered map across a range of thresholds reveals a hierarchy of structural features (Fig. 2*C*). At high thresholds, the glycan shield is composed primarily of isolated and well-defined glycans stems, while the majority of the protein surface is exposed (gray map). Moving to lower thresholds, it progresses through a series of higher-order structural states defined by isolated clusters of multiple overlapping glycans, until finally becoming completely interconnected near the noise threshold. This low threshold structure occludes large regions of protein surface and represents the full extent of the glycan shield.

As a complementary approach for visualizing the dynamic glycan shield, we performed 3-D variability analysis in the SPARX software package (27, 28) (Fig. 2*D*). 3-D variability, which is closely related to 3-D variance, should be high anywhere there is significant heterogeneity in the map. Indeed, we see high variability around known flexible regions such as the constant domains of the Fabs (clipped from view) as well as variable loops and the exterior of the protein surface at the sites of N-linked glycans. Variability in the glycans is highest at the distal ends of the glycan stems and expands outward as the threshold is reduced, revealing extensive interconnectivity (Fig. 2*D*). 2-D slices through the variability map highlight the canopy effect between neighboring glycans, which acts to occlude underlying protein surfaces (red arrows - Fig. 2*D*). Overall the shape and topology are similar to the Gaussian filtered map. A more complete description of the 3-D variability can be found in the *SI Appendix*.

### A high-throughput atomistic modeling (HT-AM) pipeline for generating large ensembles of glycosylated Env

Cryo-EM does not capture atomic details of individual molecules, so to better understand the results presented above, we performed atomistic simulations of fully glycosylated BG505 SOSIP.664. To overcome the sampling limitations of MD simulations, we employed a high throughput atomistic modeling (HT-AM) pipeline based around the ALLOSMOD package of the MODELLER software suite (29), which has been used previously to generate small ensembles of glycosylated Env (30). This methodology accounts for both energetic and spatial constraints on glycan sampling by a combination of empirical energy minimization-based structural relaxation and simulated annealing (described in detail by Guttman et al. (29), along with initial randomization of glycan orientations. However, it does not provide temporal information about dynamics. Here, we have built on this by incorporating 10 different protein scaffolds and additional variability in the V2 and V4 loops (Fig. 3*A*) and utilized it to generate much larger ensembles of 1000 fully glycosylated Env structures with uniform mannose-9 (Man9) glycans. Ten such models are shown in Fig. 3*C*, one from each protein scaffold, while the full set of models at a single glycosylation site is shown in Fig. 3*D*. We also repeated the simulation with uniform mannose-5 (Man5) glycosylation for comparison.

**Fig 3.**
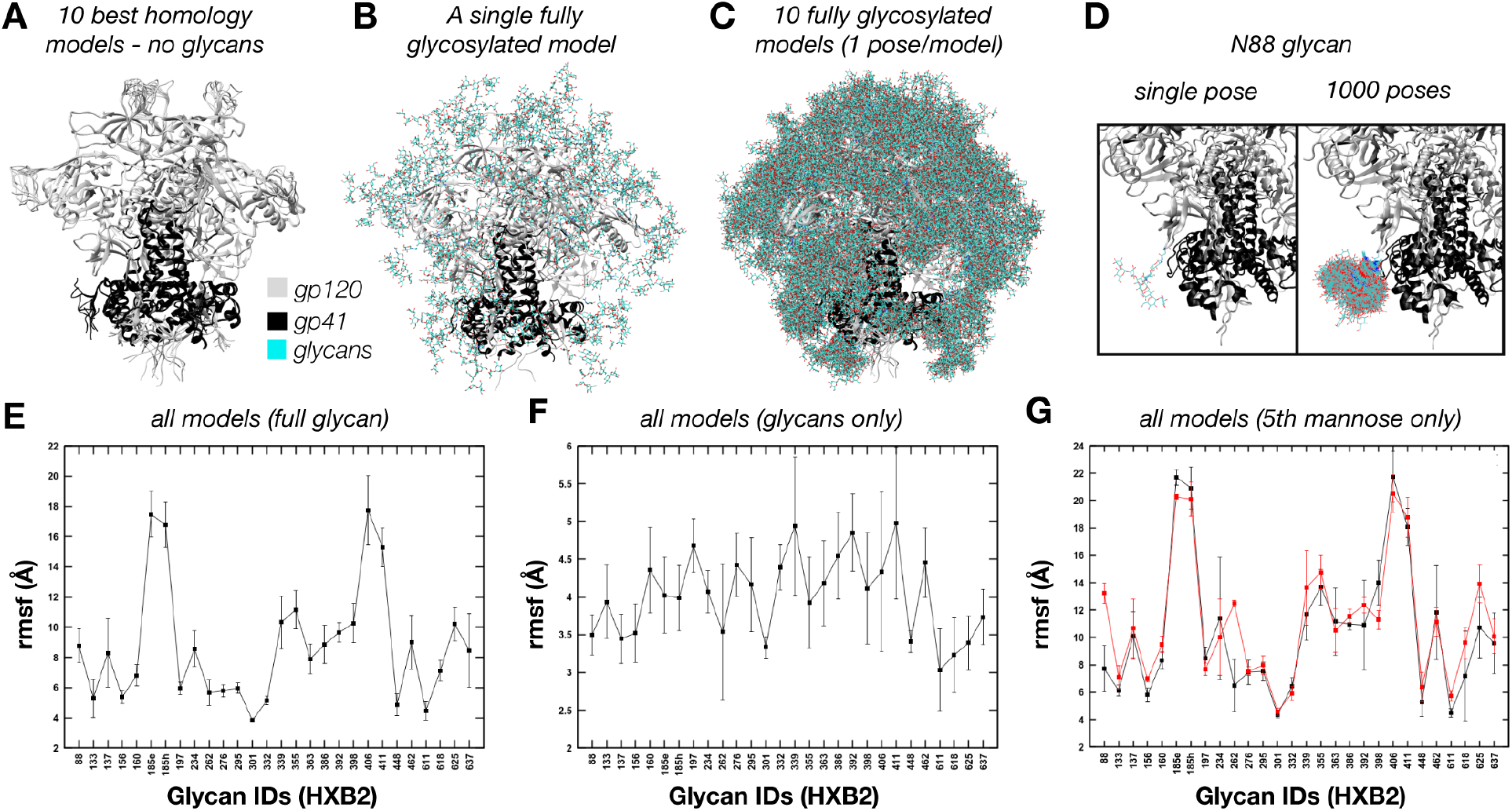
ALLOSMOD-based HT-AM pipeline for fast and robust sampling of fully glycosylated Env. **(A)** 10 homology models used as protein scaffolds. **(B)** One of the ten models with a single relaxed Man9 glycan at each site. **(C)** One fully glycosylated and relaxed model for each of the ten protein scaffolds. **(D)** Close up of the N88 glycan showing a single glycan pose (L) and all 1000 poses (R). **(E)** Full glycan average rmsf for all 1000 poses. **(F)** Glycan only rmsf for all 1000 poses. **(E)** Full glycan average rmsf for all 1000 poses at the 5^th^ mannose residue for both the Man9 and Man5 ensembles.

### The HT-AM pipeline capture spatial and energetic constraints on glycan flexibility

To assess how spatial and energetic constraints affect the flexibility of individual glycans in our simulations we calculated the root mean-squared fluctuation (rmsf) for each glycan across all 1000 models after aligning the protein scaffold. Our method captures variability between glycans (Fig. 3*E*), and the additional flexibility imparted by the 10 starting protein scaffolds can be appreciated by comparison to the glycan only rmsf (Fig. 3*F*). For example, glycans located on the flexible loops have a much higher rmsf than all other glycans (185e, 185h, 406 and 411) whereas glycans at the N262, N301, N332, N448, and N611 sites have lower rmsf. This leads to a large difference in sampled volumes between the most and least dynamic glycans (*SI Appendix*, Fig. S4*A*). We also see an increase in average rmsf of the individual glycan residues starting from the core NAG and moving outward to the tips of each antenna (*SI Appendix*, Fig. S5*A-C*), which is in line with the cryo-EM results showing reduced resolution beyond the core NAG (Fig. 1*D*).

We observed a similar trend for the Man5 ensemble (*SI Appendix*, Fig. S5*G-H*), however, by comparing the average rmsf values of the 5^th^ mannose residue alone, we found a slight increase in rmsf and sampled volume at most sites compared to the Man9 ensemble (Fig. 3*G* and *SI Appendix*, Fig. S5*I-J*). We attribute this effect to increased crowding in the glycan canopy from the more massive Man9 glycans. In support of this, there is a significant positive correlation between glycan flexibility glycan crowding (see *SI Appendix*), however, only when considering relatively large neighborhoods (*SI Appendix*, Fig. S6*A*). A similar trend was observed by Stewart-Jones et al. which they attributed to different “shells” of influencing glycans (6). Such higher-order dynamic effects could be possible in light of our structural observations.

In addition to crowding effects, the local protein structure will also influence glycan dynamics. In our modeling pipeline, the protein backbone was kept harmonically restrained close to the template to allow for extensive sampling of glycan conformations using simulated annealing. Thus, we see that the Asn sidechains of residues 88, 160, 197, 234 and 262 all have very low rmsf (*SI Appendix*, Fig. S5*F*), possibly stemming from limited torsional space available during modeling. The glycosylated Asn residues in gp41 have relatively low rmsf as well (N611, N618, N625, and N637), being situated on stable helical bundles (*SI Appendix*, Fig. S4*B*). This ultimately results in a relative reduction of the glycan dynamics at some of these sites (Fig. 3*C*). Correcting for the contribution to fluctuations coming from the underlying protein, we observed that the rmsf between the different glycans are comparable, ranging from 3 Å to 5 Å, with similar scale of SD (Fig. 3*F*).

### Simulated cryo-EM maps reproduce defining features of the experimental data

Given the single-molecule nature of cryo-EM datasets, the ensembles generated by the HT-AM pipeline can be seen as representing the individual particles that go into a 3-D reconstruction. With that in mind, we established a protocol for transforming the ensembles into simulated cryo-EM datasets (Fig. 4*A*). For comparison, we replicated the process for the Man5 ensemble as well as a protein-only ensemble (BG505_Man9, BG505_Man5 and BG505_PO respectively). The simulated maps reproduced some of the defining features of the experimental data. For instance, refinement was dominated by the stable protein core and only the first few sugar residues at each site are defined when the map is filtered to the global FSC resolution (Fig. 4*A*). This leads to a similar scale-space structure (Fig. 4*B*), with a plateau again appearing around 1.5-2 SD when the maps are filtered to the same initial resolution, suggesting that the HT-AM pipeline is capturing physiologically relevant sampling in the glycan shield, at least globally. Importantly, we found that the volume of the 1.5 SD Gaussian filtered map at the noise threshold closely approximates the total volume sampled by the ensemble (Fig. 4*B* - dashed lines). Comparing the curves for all three simulated reconstructions (Fig. 4*B*) we see how differences in glycosylation manifest as differences in volume at low resolutions, suggesting cryo-EM can be used to capture global changes in glycan shield composition between two reconstructions of the same Env. Visualization of the 1.5 SD Gaussian filtered BG505_Man9 map at high and low thresholds reveals a similar evolution towards a more connected topology as the threshold is reduced and extensive shielding of protein surface (Fig. 4*C*). We also calculated the 3-D variability map for the BG505_Man9 reconstruction and observed a similar result to the experimental data (Fig. 4*D*). In addition, we compared the SPARX variability map to the true 3-D variance (*SI Appendix*, Fig. S7*B*) and found negligible differences between the two (See the *SI Appendix* for details).

**Fig 4.**
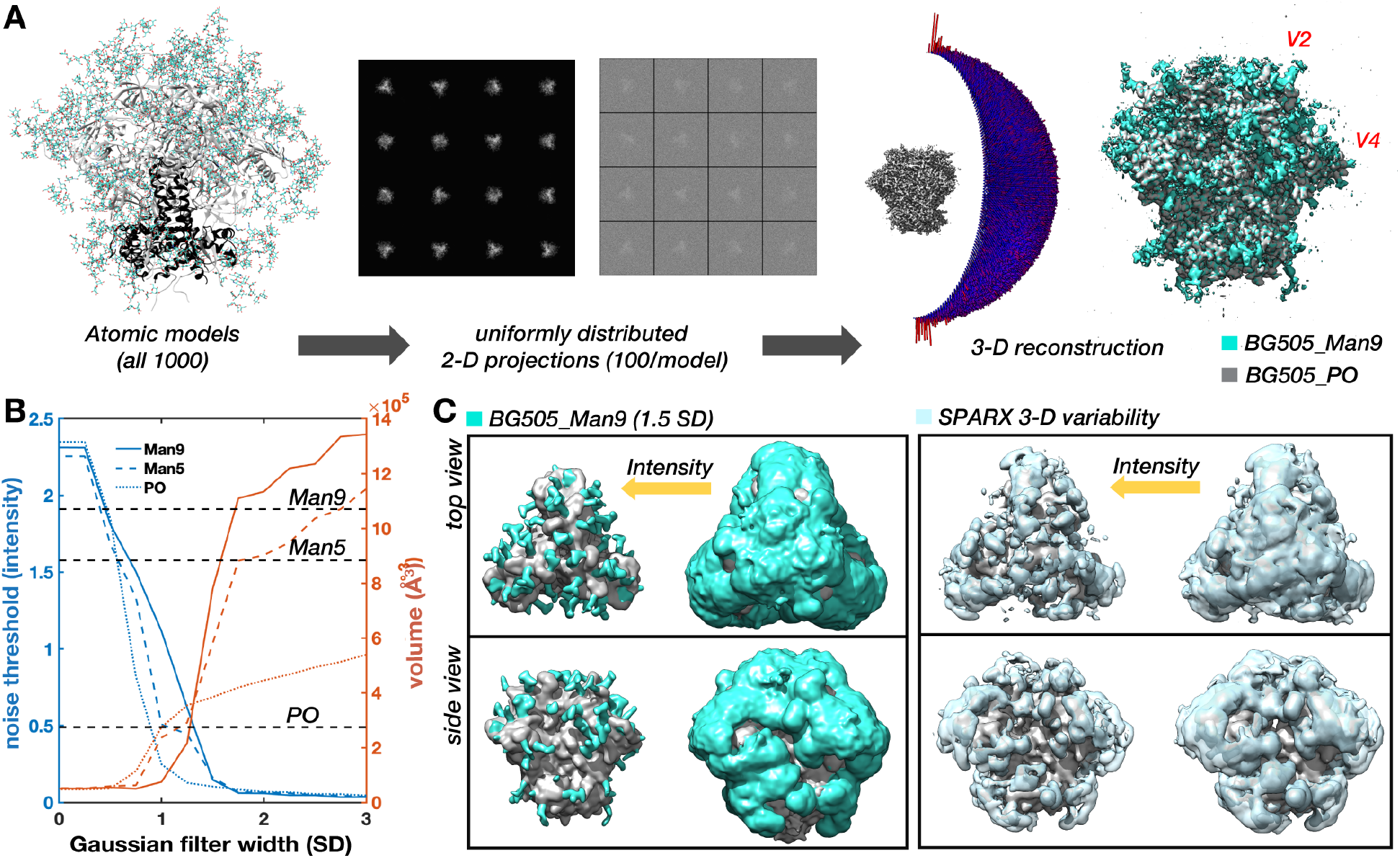
Simulated cryo-EM maps from the HT-AM ensembles reproduce defining features of the experimental data. **(A)** Schematic of the simulated data creation pipeline. Each model is projected at 100 uniformly distributed angles (representative 2-D projections with and without noise are shown), then the combined dataset of 100K projections is refined using RELION (the resulting 3-D reconstruction and angular distribution histogram are shown). **(B)** Noise threshold and volume at the noise threshold as a function of Gaussian filter width (SD) for the BG505_Man9, BG505_Man5, and BG505_PO simulated cryo-EM maps with dashed lines indicating the volume sampled by each ensemble. **(C)** 1.5 SD Gaussian filtered BG505_Man9 (teal) and BG505_PO (gray) maps along with the SPARX 3-D variability map (light blue) viewed from the side and top at high and low ntensity thresholds.

### Measuring glycan dynamics in cryo-EM maps

Measuring intensity around individual glycans in the cryo-EM maps should allow us to assess their relative dynamics. To confirm this, we used the simulated BG505_Man9 map since we could make direct comparison to the rmsf values. However, only the first 1-2 glycan residues are defined at most sites in the high-resolution map (Figure5A) and their dynamics deviate significantly from the rest of the glycan (*SI Appendix*, Fig. S8*A*). The third glycan residue, β-mannose (BMA), more closely approximates the average rmsf (mean deviation ~2Å) and the relative differences between sites (*SI Appendix*, Fig. S8*A-B*). In addition, the glycan stems are well defined in the 1.5 SD Gaussian filtered map, which allowed us to identify their locations in the map with reasonable accuracy. So, we built and relaxed glycan stems into the 1.5 SD Gaussian filtered BG505_Man9 simulated map (Fig. 5*A*) at every glycosylation site we could confidently identify (23/28 per protomer). We could not identify the V2 and V4 loop glycans at N185e, N185h, N406, and N411, which had the highest rmsf values in the simulation, nor the glycan at N339, which has a lower rmsf but projects directly towards the heterogeneous V4 loop. The positions of the BMA residues were then used to analyze local map statistics within a spherical probe around each glycan (Fig. 5*A*). We observe a strong positive correlation between the inverse rmsf and normalized mean intensity (Fig. 5*B-C*), with the peak correlation coefficient of ~0.88 (p=2.1e-8) occurring when using the unfiltered map and a probe radius of 2.3Å. Thus, local intensity around BMA residues accurately captures ground-truth differences in relative dynamics between glycans and can be used to make direct comparisons between the simulated and experimental cryo-EM maps.

**Fig 5.**
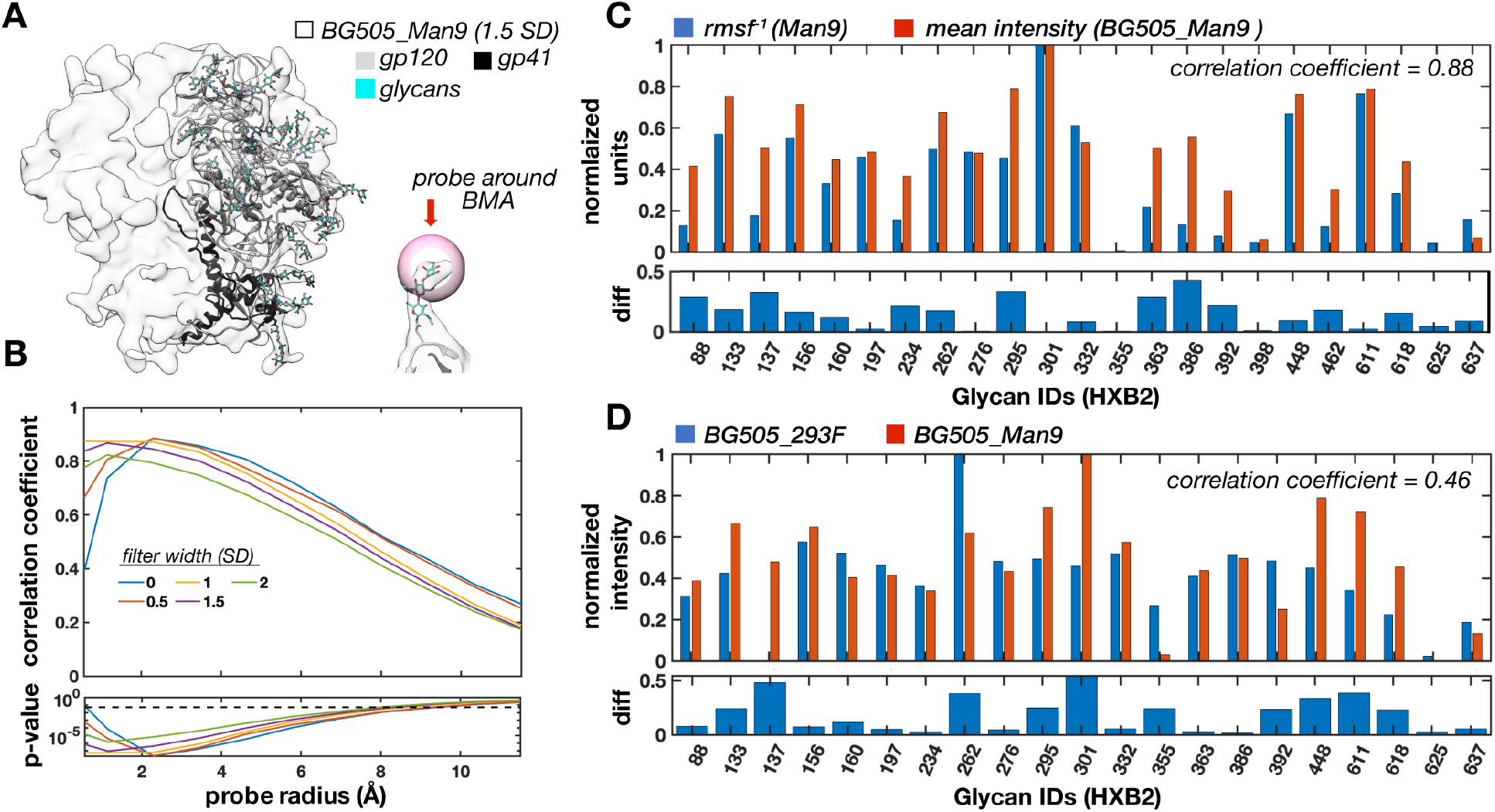
Measuring glycan dynamics from cryo-EM maps reveals close agreement between simulation and experiment. **(A)** 1.5 SD Gaussian filtered BG505_Man9 simulated cryo-EM map (transparent) and atomic model used for local intensity analysis. Also shown is a single glycan stem with a spherical probe around the BMA residue. Pearson correlation coefficient and p-value between mean intensity and inverse rmsf as a function of probe radius for 5 different Gaussian filtered maps. The dashed line designates the confidence threshold (p-value = 0.05). Normalized mean intensity around BMA residues for BG505_Man9 (probe radius = 2.3Å) and the inverse normalized full glycan average rmsf. Pearson correlation coefficient ~0.88 (p-value = 2.1e-8). (D) Normalized mean intensity around BMA residues for BG505_293F and BG505_Man9 along with the absolute difference at each site. Pearson correlation coefficient ~0.46 (p = 0.03).

### The HT-AM pipeline reproduces physiologically relevant trends in glycan dynamics measured by cryo-EM

With a method in place for measuring glycan dynamics from cryo-EM maps, we then made direct comparisons with the experimental data. We built and relaxed glycan stems into the 1.5 SD Gaussian filtered BG505_293F map as described above and could identify clear density at 21/28 glycosylation sites per monomer, two less than from the simulated map (N398 and N426). The other five missing glycans were common to both maps, meaning the HT-AM pipeline captured physiologically relevant dynamics at these sites, at least up to the detection limits of this method. Overall, we found that the HT-AM pipeline captured a similar trend in glycan dynamics with a correlation coefficient between the two of ~0.46 (*p* = 0.03) (Fig. 5*D*).

In addition to the V4 and V5 loop glycans at N398 and N462, we also saw a large deviation at the N137 glycan on the V1 loop, as well as the N262 and N301 glycans. In the BG505_293F map, the N262 glycan was the most ordered due to stabilizing contacts with the gp120 core, and these interactions may not have been accurately captured by the simulation given the restricted protein dynamics. In gp41, a large deviation also occurred at the N611 glycan, which we attribute to restrictive sampling of the protein backbone as previously discussed. To complicate the comparison, the N618 and N625 sites are under-occupied as revealed by MS (*SI Appendix*, Fig. S2*A*). Although sub-occupancy will cause reduced signal intensity due to averaging, the severity of this effect is diminished when dynamics at that site are large relative to other glycans, which is the case for the glycan at N625.

### Detecting site-specific changes in glycan dynamics, occupancy and chemical composition from cryo-EM maps

To test whether these methods could be used to detect site-specific changes in dynamics and occupancy between cryo-EM maps of differentially glycosylated Env, we performed a comparative analysis between the BG505_Man9 and BG505_Man5 simulated maps. Given that stems of a Man9 and Man5 glycan are identical, the only changes in intensity around the BMA residue should arise from differences in dynamics alone. On average, we see a ~17% reduction in intensity indicative of increased dynamics, which is in line the rmsf data (Fig. 6*A*). We also accurately detect the largest increase and decrease in dynamics at the N262 and N234 sites respectively. To verify we could detect changes in occupancy, we removed the glycan at the N625 site from half of the models and re-refined the data (referred to as BG505_Man9HO for “half occupancy”). Not surprisingly, we see a ~50% reduction in mean intensity from the fully occupied reconstruction around this site (Fig. 6*A*).

**Fig 6.**
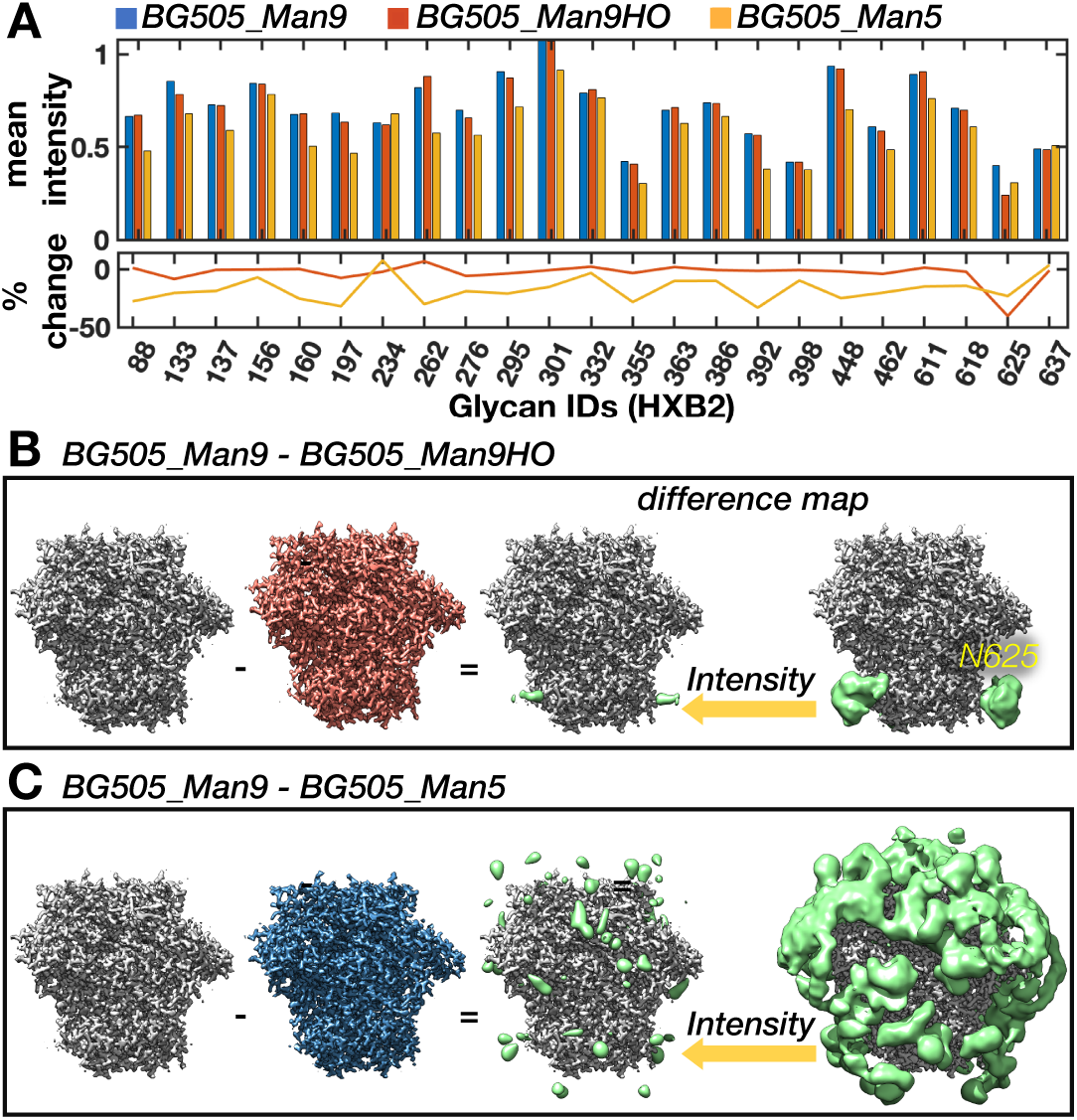
Detecting changes in glycan dynamics, occupancy, and chemical composition from simulated cryo-EM maps. **(A)** Mean intensity around each glycan BMA residue for the BG505_Man9, BG505_Man5, and BG505_Man9HO maps along with the percent change from BG505_Man9. **(B)** BG505_Man9 - BG505_Man9HO and (C) BG505_Man9 - BG505_Man5 difference maps (Gaussian filtered) at two intensity thresholds (right - green) along with high-resolution sharpened maps (left).

Another technique that should be sensitive to subtle changes between two cryo-EM maps is difference mapping. Indeed, the change in occupancy at the N625 site is apparent in the BG505_Man9 - BG505_Man9HO difference map (Fig. 6*B*). At high threshold, the signal is localized around the glycan stem and extends to the protein surface. In the BG505_Man9 - BG505_Man5 difference map (Fig. 6*C*), however, the difference signal is strongest where the distal tips of the Man9 glycans would be and does not extend to the protein surface. These results establish cryo-EM as a tool for measuring glycan dynamics as well as changes in chemical composition and occupancy between differentially glycosylated Envs.

### Insights gained from analysis of simulated data allows improved characterization of cell-type specific differences in glycan shield

To test our methods experimentally, we collected cryo-EM data on complexes of RM20A3 Fab and BG505 SOSIP.664 expressed in two additional common cell lines that produce major and minor changes in glycosylation; HEK293S cells, and a stable CHO cell line (referred to as BG505_293S and BG505_CHO respectively). The stable CHO cell line expressed Env sample was provided to us by IAVI as part of the demo run conducted prior to the currently on-going human clinical trials, and should be identical to the final vaccine product (31). Both new datasets refined to ~3Å-resolution (*SI Appendix*, Fig. S1*A*) and the three atomic structures are nearly identical (Cα RMSDs ~ 1Å - *SI Appendix*, Fig. S1*B*). Scale-space (*SI Appendix*, Fig. S3*G-I*) and 3-D variability analysis (*SI Appendix*, Fig. S9*A-C*) also revealed similar low-resolution structural features to BG505_293F, however, comparison of local map intensity around BMA residues uncovered some significant difference in the gp41 glycans, particularly at the N611 and N625 sites, potentially from changes in occupancy (Fig. 7*A*). This is further supported by site-specific MS data collected for the two additional samples (*SI Appendix*, Fig. S2*A*).

**Fig 7.**
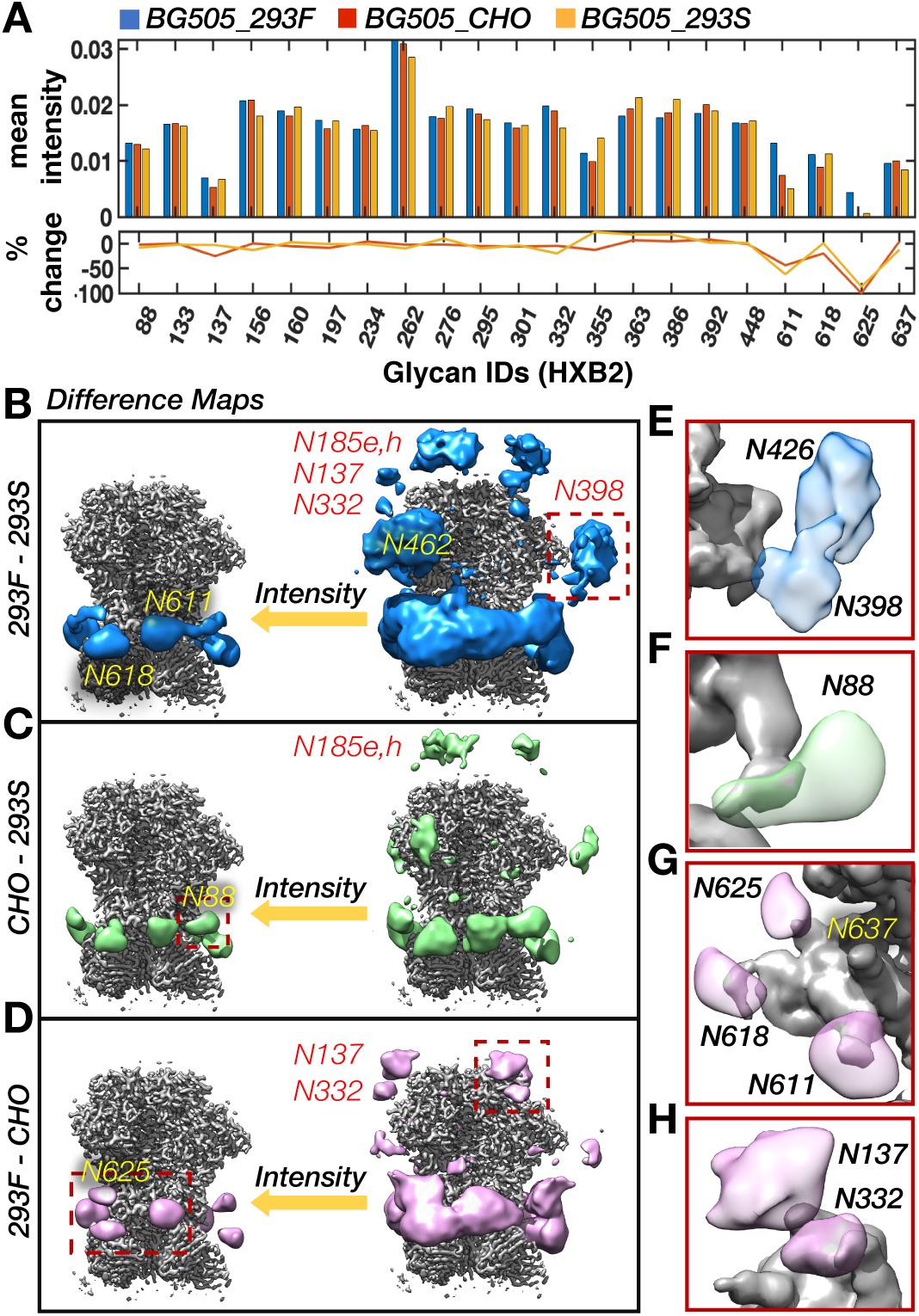
Detection of cell-type specific changes in glycan shield composition and dynamics with cryo-EM. (**A**) Mean intensity at each glycan BMA residue for the BG505_293F, BG505_293S, and BG505_CHO cryo-EM maps along with percent change from BG505_293F. **(B)** BG505_293F – BG505_293S, **(C)** BG505_CHO – BG505_293S, and BG505_293F - BG505_CHO difference maps at two intensity thresholds. All difference maps were multiplied by a soft mask around the RM20A3 Fabs then smoothed with a 2 SD Gaussian filter. **(E-H)** Fly-outs of the regions outlined by dashed red boxes.

The 293S sample contains only high-mannose type glycans and should therefore allow detection of complex glycans via difference mapping, and to a lesser extent, different distributions of high-mannose glycans. Indeed, we see strong difference signal around the primarily complex gp41 glycans in both the difference maps (Fig. 7*B-C*). We also observe signal around the primarily complex glycan at N88 (Fig. 7*F*). In addition, clear difference signal appeared around the V2, V4, and V5 loops, specifically near the glycans at N185e and h, N398, and N462, all of which are shown to be complex by MS. The MS data also shows a subtle difference in both occupancy and percentage of complex glycans at the N398 site between the 293F and CHO sample, which could explain why there is strong difference signal in one map and not the other. According to the MS data, the only significant differences in occupancy between the 293F and 293S samples occurred at the N137, N133, and N611 sites, however the difference maps do not contain any signal around the N611 glycan, suggesting there may be discrepancies between the two methods.

The MS analysis also detected differences in occupancy between the CHO sample compared to the 293F sample at multiple sites (*SI Appendix*, Fig. S2*A*). Indeed, upon closer examination we found that the difference signal around the gp41 glycans at N611, N618, and N625 extends all the way to the protein surface (Fig. 7*G*), indicative of changes in occupancy. In this difference map, there is also clear signal around the N137 site, and to a lesser extent at the tip of the N332 glycan stem (Fig. 7*H*). Given the proximity of N137 and N332, it is plausible that sub-occupancy at one is driving changes in dynamics and/or glycan distribution at the other. Similar higher-order effects have also been observed via MS when comparing glycoform distributions on Env before and after knocking out specific glycans (26, 32, 33).

### A probabilistic glycan-glycan interaction network reveals highly connected glycan clusters

Both the experimental and simulated cryo-EM maps showed extensive interconnectivity among glycans, so we sought to quantify this more precisely. With each glycan in our models sampling a particular region of space, neighboring glycans can explore overlapping volumes, and the fraction of this overlap gives a measure of their interaction probability. Fig. 8*A* shows a heat map of the normalized glycan-glycan volume overlap matrix. By interpreting this as an adjacency matrix we can model the glycan shield as a network. Here, each glycan was defined as a node, and two nodes were connected by an edge if there was at least 5% overlap between them (Fig. 8*B*). The only glycans from the neighboring protomers that were considered are those with an inter-protomer edge. Fig. 8*C* shows the network obtained for the Man9 ensemble unfolded in 2-D. It can be seen that overall there are three main regions of overlap, – the V1/V2 apex, the gp41 base, and the densely occupied gp120 outer and inner domains that includes the high-mannose patch (HMP). The inter-protomer overlaps are contributed mainly by the V1/V2 glycans. The nomenclature was inspired by the established nomenclature for gp120 domains (34), however, our definitions were adapted to better capture glycan shield structure (*SI Appendix*, Fig. S10*A*). The analysis was repeated for the Man5 ensemble and we observed a reduced overall connectivity as measured by the mean node degree (from ~7 to ~5) and the maximum network diameter (from 5 to 8 hops), consistent with their smaller size (*SI Appendix*, Fig. S*11*).

**Fig 8.**
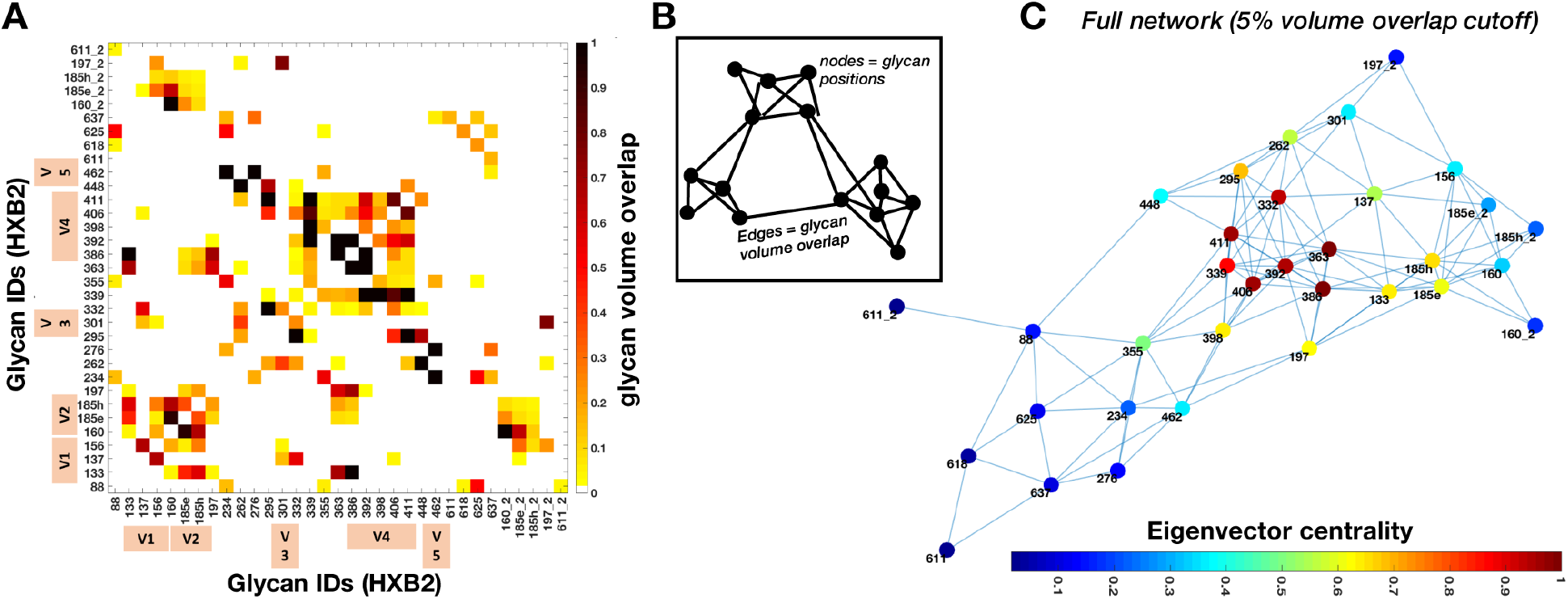
Probabilistic glycan-glycan interaction network for the Man9 ensemble. **(A)** Glycan-glycan volume overlap matrix. Inter-protomer overlap is given by suffix “_2” and the variable loop glycans are indicated by tan color bars. The overlap fraction is normalized with at least 50% overlap being designated as 1. **(B)** Cartoon model of the glycan-glycan interaction network generated by interpreting the overlap matrix as an adjacency matrix. The edge length is drawn inversely proportional to the overlap value. Edges represent overlap between two glycans represented by nodes. **(C)** Glycan-glycan interaction network calculated from the matrix in (A) with nodes colored by normalized eigenvector centrality.

With a network in place we could then analyze the relative influence of each glycan on the whole system and examine its long-range structure. To do this we calculated the relative eigenvector centrality of the nodes, which is a measure of importance in the network, and the results are projected on the network as a colormap in Fig. 8*C*. We see that the gp120 outer domain glycans at N332, N339, N363, N386, and N392, which are densely connected within the network, have strong eigencentrality measures. In the apex region, the V1/V2 loop glycans at N133, N160 and N185e/h also have relatively high eigencentrality due to their increased flexibility, while the glycans at N88, N234, N276, and those in gp41 have low eigencentrality, reflecting low interaction probabilities. Incorporated intrinsically into the network is a set of stable sub-graphs that represent highly connected glycan clusters. To illustrate this structural hierarchy, we progressively stripped the network using tighter overlap cutoffs (*SI Appendix*, Fig. S10*B*). As the network is degraded, we see the formation of two large sub-graphs; one composed of the V1/V2 apex and the gp120 outer domains; and the second composed of the gp120 inner domain along with gp41 base glycans. With an even stricter cutoff, the sparsely connected glycans with low eigencentrality separate out and the two sub-graphs split again to form four sub-graphs; a V1/V2 apex domain; a gp120 outer domain; a gp120 inner domain; and lastly, the group of unconnected gp41 base glycans.

### Highly connected glycan clusters are resistant to enzymatic digestion and help stabilize of the pre-fusion trimer

To determine if the glycan shield affects Env structure and dynamics we exposed BG505_293S (already in complex with RM20A3 Fab) to digestion by endoglycosidase H (Endo H), which cleaves high-mannose glycans between the first and second sugar rings (Fig. 9*A*), and collected cryo-EM data on the deglycosylated sample. In addition, we hypothesized that highly connected glycans would be protected from enzymatic digestion, and vice versa. Therefore, if we could measure the order in which glycans were cleaved by Endo H it could provide indirect validation of our network models. To test this, we collected cryo-EM data on a partially deglycosylated sample as well. The reaction times and Endo H concentrations for achieving partial (2hrs) and complete (16hrs) digestion were determined prior to performing cryo-EM. The datasets, referred to as BG505_EndoH2 and BG505_EndoH16, reconstructed to ~3.2Å and ~3.5Å-resolution respectively, with similar overall quality, resulting in nearly identical atomic models to the other three datasets (*SI Appendix*, Fig. S1*A-B*). The progressive digestion of the glycan shield was confirmed by SDS-PAGE and SEC chromatography (*SI Appendix*, Fig. S12), while scale-space analysis of the cryo-EM maps revealed a clear progressive reduction of glycan signal (*SI Appendix*, Fig. S3*G-I*).

**Fig 9.**
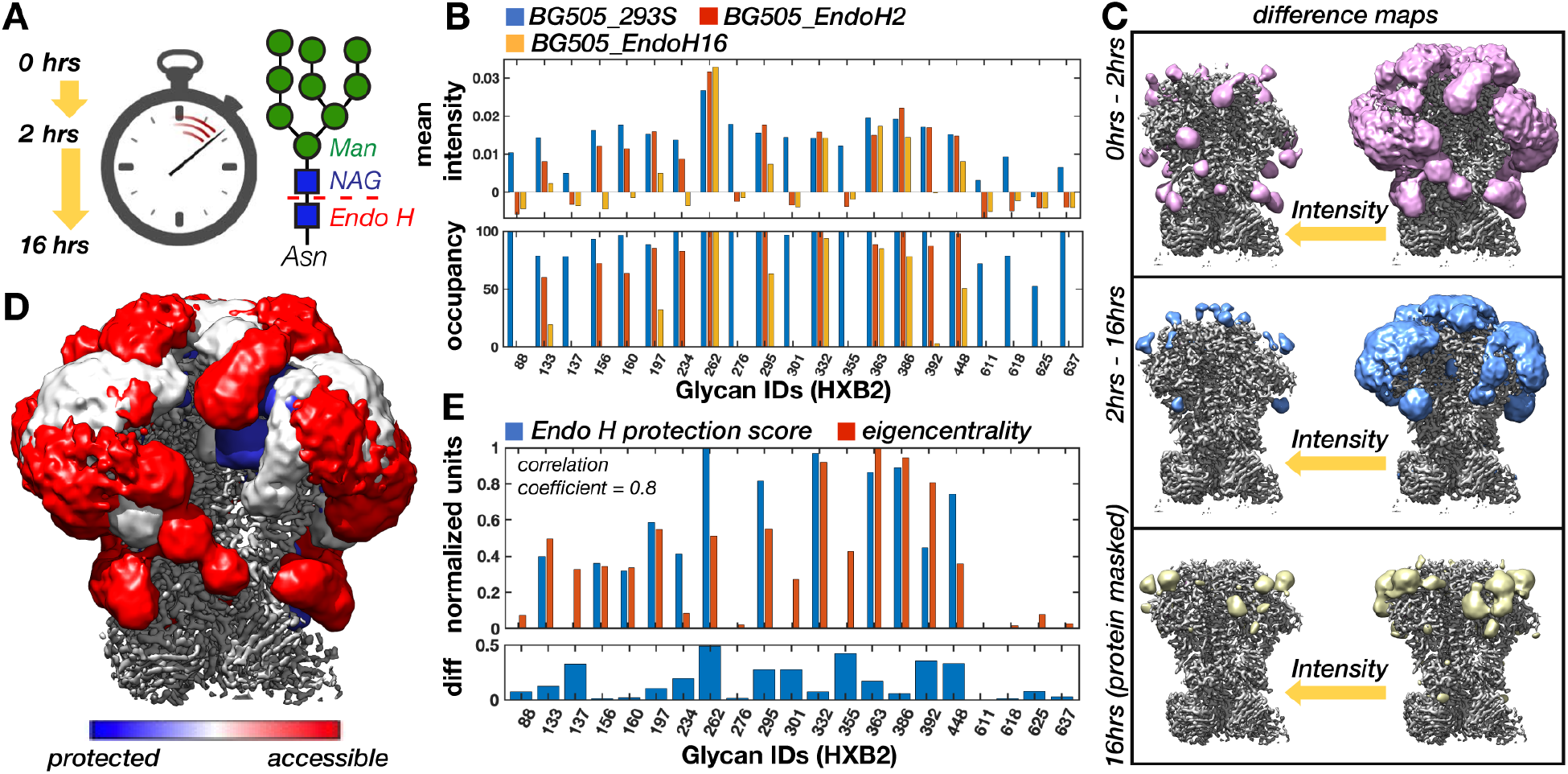
Time-resolved cryo-EM reveals that highly connected glycan clusters are resistant to enzymatic digestion. **(A)** Cartoon schematic of the Endo H digestion of high-mannose glycans and an illustration of the three reaction times captured by time-resolved cryo-EM. **(B)**Mean intensity around glycan BMA residues for the BG505_293S, BG505_EndoH2, and BG505_EndoH16 cryo-EM maps along with the percent occupancy at each site. Occupancy was calculated assuming a linear relationship between occupancy and intensity with the initial occupancy determined by MS. (C) BG505_293S – BG505_EndoH2 and BG505_EndoH2 – BG505_EndoH16 difference maps at two intensity thresholds. Difference maps were first multiplied by a soft mask around the RM20A3 Fabs and smoothed with a 2 SD Gaussian filter. Also shown is the residual glycan signal remaining after 16hrs of digestion isolated by masking out the protein density. **(D)** Overlay of the three maps from panel B colored according to susceptibility to Endo H digestion. **(E)** Normalized Endo H protection score measured as the cumulative occupancy at each site in the two digestion intermediates and the normalized eigencentrality from the Man9 volume overlap network. Pearson correlation coefficient of ~0.8 (p=1.14e-05).

Indeed, analysis of local map intensity around each glycan revealed that digestion occurred non-uniformly (Fig. 9*B-C*). This was also confirmed by changes in the 3-D variability maps (*SI Appendix*, Fig. S9*D-E*). By assuming a linear relationship between intensity and occupancy and using the MS data to set the initial occupancy, we calculated the occupancy at each site (Fig. 9*C*). After 2 hours the gp41 glycans (N611-637) were completely digested, while some glycans, particularly those in the densely packed gp120 outer domain, remained mostly intact. Although we could not identify unique signal for the individual V2, V4, and V5 loop glycans, there was clear signal around these sites in the 0-2hr difference map only (Fig. 9*B*), indicating the dynamic V-loop glycans were completely digested within 2hrs. We also found partial signal reduction at a few sites, indicative of non-uniform digestion. For example, the apex cluster composed of the N156 and N160 glycans as well as the glycans at N133, N197, and N234. After 16 hours the glycan shield was almost completely digested, however, we still detected some residual glycan signal at the previously discussed cluster composed of the N363, N386, and N197 glycans, as well as a cluster composed of the N295, N332, and N448 glycans. In addition, the highly protected glycan at N262 remained completely intact. By quantifying the degree of protection from Endo H (see Methods) and comparing it to the predicted network eigencentralities for the Man9 ensemble (Fig. 9*E*), we obtained a correlation coefficient of ~0.8 (p=1.14e-05), suggesting highly connected glycans are resistant to enzymatic digestion. Also evident is the similarity between the persistent glycan clusters and the sub-graphs generated by progressively stripping the glycan overlap network (*SI Appendix*, Fig. S10*B*), which could be seen as mimicking the gradual digestion by Endo H.

Interestingly, 3-D classification of the Endo H treated datasets revealed an increasing degree of protein unfolding and subunit dissociation initiating at the V1-3 loops in the trimer apex. Classification consistently converged to four 4 classes that appeared to capture found heterogeneous intermediates in the unfolding process (Fig. 10*A* and *SI Appendix*, Fig. S13-17). We interpreted these as representing (1) stable trimers, (2) trimers with unfolded or partially unfolded V1-V3 loops, (3) trimers with fully unfolded/dissociated gp120s, and (4) monomers/dimers. As the reaction progressed, the percentages of unfolded trimers increased, while the percentage of stably folded trimers decreased (Fig. 10*B*). Although it cannot be easily confirmed, the unfolded trimers in each dataset were likely more completely deglycosylated than the particles that make up the stable trimeric classes. We also observed a small percentage of dissociated gp120s in the CHO sample, perhaps as a result of the higher degree of sub-occupancy. When viewed in context of the results presented above, we concluded that the highly connected glycans and glycan clusters that were resistant to digestion are also critical to maintaining structural stability of the pre-fusion Env trimer, suggesting the glycan shield may have additional functions beyond its role in immune evasion.

**Fig 10.**
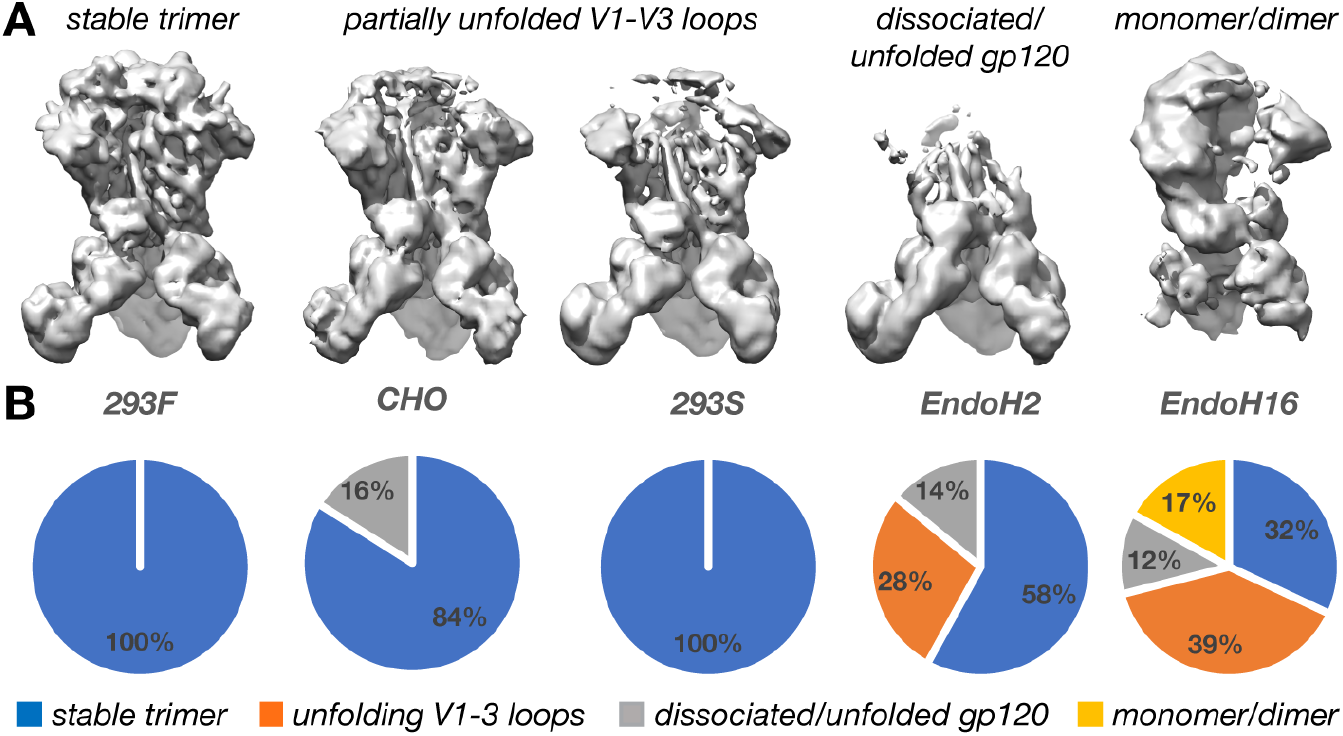
Digestion of the glycan shield leads to progressive destabilization of the Env timer. **(A)** Representative cryo-EM maps of the unfolding intermediates captured by 3-D classification. **(B)** Pie charts showing the relative percentages of each class present in the five cryo-EM datasets presented here.

## Discussion

Prior to this study, there were conflicting reports of glycan shield structure from cryo-EM and X-ray crystallography measurements (6, 14, 35). Our results clearly show that the native glycan shield is highly dynamic with respect to the protein core. We did not observe any stabilized glycan-glycan interactions like those reported in crystal structures (6), suggesting methodological differences may be responsible for the apparently contradictory results. For example, the crystal structures included two bound bnAbs per monomer to facilitate crystal packing, both of which engage multiple glycans and disrupt the native dynamics and higher-order structure. Desolvation of mobile waters embedded within the glycan shield during crystallization could also potentially induce the stabilized glycan-glycan contacts observed. A favorable interpretation that also fits with the conclusions of this paper is that the crystal structures captured artificially stabilized yet physiologically relevant conformations and are thus suggestive of the types transient, of non-covalent interactions possible between glycans.

Our work represents the first case of using cryo-EM to validate glycoprotein ensembles, and in turn, their use in guiding cryo-EM analysis. We found that the HT-AM pipeline captured physiologically relevant glycan sampling at global and local scales, however, deviated from the experimental observations when the flexibility of the underlying protein was greater than could be captured by the current pipeline. Thus, exploring methods to enhance sampling of the protein backbone, particularly at Env variable loops, represents a clear route for improving the accuracy of the pipeline (36). Another logical next step is to use the experimental cryo-EM maps to steer the ensemble modeling process, which has been successful for non-glycosylated proteins (37–39), however, extending such techniques to capture the extreme levels of heterogeneity present in the glycan shield will be challenging. Experimentally, our ability to capture differences in glycan dynamics from cryo-EM maps is limited when using local map intensity around BMA residues alone, and such limitations could also be contributing to the observed discrepancies. Again, using the cryo-EM maps to steer the modeling process could provide more accurate and comprehensive assessments of variations in relative dynamics between glycan in addition to improving modeling accuracy. Finally, utilizing sequence engineering and/or different expression systems to achieve a more uniformly glycosylated Env sample, or conversely, incorporating experimentally relevant levels of heterogeneity into the modeling pipeline will allow for a more accurate comparison between theory and experiment.

Using graph theory, we mapped out the complex network of potential interactions within the glycan shield. A similar network-based approach was introduced previously to capture concerted behavior within the glycan shield during all-atom MD simulations (22). Although the HT-AM pipeline does not capture temporal dynamics, we found that the probabilistic networks reported here closely resembled the ones derived from the MD simulations, and the two lead to similar predictions. For instance, the clustering of glycans into larger microdomains (*SI Appendix*, Fig. S10*A-B*). Moreover, the overlap probability between glycans gives a measure of the structural shielding rather than the energetic interactions between glycans. At a qualitative level, the hierarchy of structural features we observed in the Gaussian filtered and 3-D variability maps (Fig. 2*C-D*) could be seen as the ensemble-average manifestation of this clustering behavior. Given the preference of bnAbs for targeting the interfaces between higher-order glycan clusters (22), experimentally mapping such long-range structure could be important. At a more quantitative level, we found that the average degree of protection from Endo H digestion correlated strongly with network eigencentrality for those glycans we could identify in the cryo-EM maps, and that the order of digestion closely matched the order in which glycans were removed from the network when applying stricter overlap thresholds. It should be noted, however, that because of the somewhat simplified approach we have taken to modeling the most dynamic segments of the variable loops, in particular in the V4 loop segment containing the N398, N406, and N411 glycans, the eigencentrality of these glycans may be artificially high, leading to minor discrepancies between digestion susceptibility and network centrality. Nonetheless, these correlations could be seen as providing indirect validation of the network models, where the high centrality glycans are either located in densely packed regions where Endo H would have a difficult time accessing (*SI Appendix*, Fig. S6*D-E*) or are protected by strong interaction probability with their neighbors.

We demonstrated that cryo-EM is capable of detecting and quantifying changes in glycan dynamics, as well as occupancy and chemical composition between differentially glycosylated Envs, the latter of which were exclusively provided by MS prior to this study. Thus, cryo-EM provides validation for these measurements while contributing insight into the structural impact of changes in glycosylation. For instance, semi-quantitative methods to assess the distribution of unoccupied sites have only recently become available (13), and here we showed that cryo-EM is highly sensitive to occupancy. Importantly, we found evidence for more extensive changes in glycosylation between the 293F and CHO expressed samples than detected by MS, which has clear relevance to the currently on-going human clinical trials based on the CHO expressed Env. In addition to these results, we found that the extent of processing measured by the percentage of high-mannose type glycans at each site was strongly correlated with ordering in the cryo-EM map (*SI Appendix*, Fig. S2*C-D*), suggesting it may be possible to predict the degree of processing at each site from cryo-EM maps alone. The relationship between these variables likely reflects their mutual dependence upon glycan crowding (*SI Appendix*, Fig. S6*B-C*), as well as the properties surrounding protein structure, both of which can reduce dynamics and restrict the access of Endo H and glycan processing enzymes in the cell.

The observation that enzymatic deglycosylation leads to progressive destabilization of the Env trimer was somewhat surprising because recent investigations into the effect of glycan knockouts and deglycosylation on Env stability and viral infectivity have come to different conclusions (40–44). However, there is ample biophysical evidence that glycans influence protein stability, dynamics, and folding, even in the case of Env (17, 45–56), although it is not obvious which mechanisms are important here. One possible mechanism is stabilizing interactions between the core NAG and neighboring side chains, which are observed throughout Env and are common in other glycoproteins. However, Endo H leaves the core NAG attached, so the stabilizing effect must arise by other mechanisms. It has been proposed that glycans stabilize proteins primarily by destabilizing the unfolded state and by increasing the height of the unfolding energy barrier (47). The same study also found that glycosylation reduced the entropy of the folded state, possibly by imposing pressure on the protein and dampening dynamics, similar to the effects of molecular crowding or confinement, although the effect was canceled out by an equivalent change in enthalpy. Along these lines, the slightly lower resolution of the BG505_EndoH16 map compared the fully glycosylated maps could be indicative of increased dynamics. In addition, our observation that the Man9 glycans were less dynamic than the smaller Man5 glycans in our simulations could be attributed to more crowding in the canopy, which in turn could further stabilize the underlying protein. Also, the stabilizing effects were found to depend strongly on the number of glycans as well as their positions on the protein (47), which is in line with the results of our Endo H digestion experiment. Therefore, given the high density of glycans on the surface of Env and the importance of their tertiary arrangement into high density clusters, it is plausible these stabilizing effects could be amplified. Furthermore, glycan-glycan interactions such as those observed in the crystal structures solved by Stewart-Jones et. al. (6) could also play a role in stabilizing folded state. Although our results clearly show that such interactions must be more dynamic than the crystal structures suggest, the combined effect of many such short-lived interactions could become significant.

The time-resolved Endo H digestion experiment provide an explicit mapping of the importance of each glycan toward maintaining the integrity of the glycan shield and the overall stability of Env. In addition to the glycan at N262, the two highly connected clusters in the gp120 outer domain centered around the N386 and N295 glycans were the most important for trimer stability, and to a lesser extent the apex cluster composed primarily of the N156 and N160 glycans. In support of this, previous studies have shown that knocking out the glycans at N386 or N262 individually have dramatic effects on trimer stability (33), whereas deletion of up to 5 glycans at once peripheral to the CD4bs, which are not part of these cluster, are well tolerated (10, 57, 58). In addition, the glycans at N386 and N295 in particular were found to strongly control the processing of the other glycans near them, reinforcing their importance in maintaining the integrity of large-scale glycan shield structure (33). These results have implications for rational vaccine design where removing certain glycans could have undesired effects on protein stability and suggest that there are limits to the number glycan deletions that can be tolerated, with particular emphasis on maintaining the integrity of the aforementioned clusters. In addition, we found that instability originated at the trimer apex, around the V1-3 loops. This region of the structure is known to be metastable and to change conformation upon CD4 and co-receptor binding(34, 59). Therefore, it is plausible the glycans and glycan clusters in this region are contributing to maintaining Env in a metastable state poised for receptor binding. Given the high conservation of the glycan clusters identified here, our results are likely universal to all Env sequences. Lastly, it is possible that glycan induced stabilization of Env may enable hyper-mutation of the underlying protein surface, thereby endowing Env with the ability to escape immune pressure with otherwise deleterious mutations.

It is known that immune responses to Env preferentially target glycan-depleted surface area (9, 10), and the results presented here provide the first experimentally determined mapping of this surface. Under reasonable physical assumptions, the ensemble-averaged structure captured by cryo-EM should be identical to the time-averaged structure of a single thermally fluctuating Env molecule at the temperature prior to vitrification. The dynamics of the glycans create a cloud/shield over the protein, thus an approaching antibody with relatively torpid dynamics will effectively “see” this blurry ensemble-average structure, which is consistent with the relatively slow on-rates of most HIV bnAbs. We quantified the shielding effect with a simple “rolling sphere” method for BG505_CHO and visualized the results as a colormap projected onto the surface of Env (Fig. 11). White surfaces are strongly masked by nearby glycans and red surfaces are weakly masked, highlighting sites of potential vulnerability to neutralizing antibodies (see the *SI Appendix* for details). Although the cryo-EM maps presented here do not perfectly capture the in-vivo structure of a single fluctuating Env because they are ensemble averages of chemically heterogeneous molecules, if one considers the net serum response to a vaccine that itself contains the exact same heterogeneity as the sample we analyzed here (60, 61), they still provide an accurate representation of the average surface exposure of the entire ensemble.

**Fig 11.**
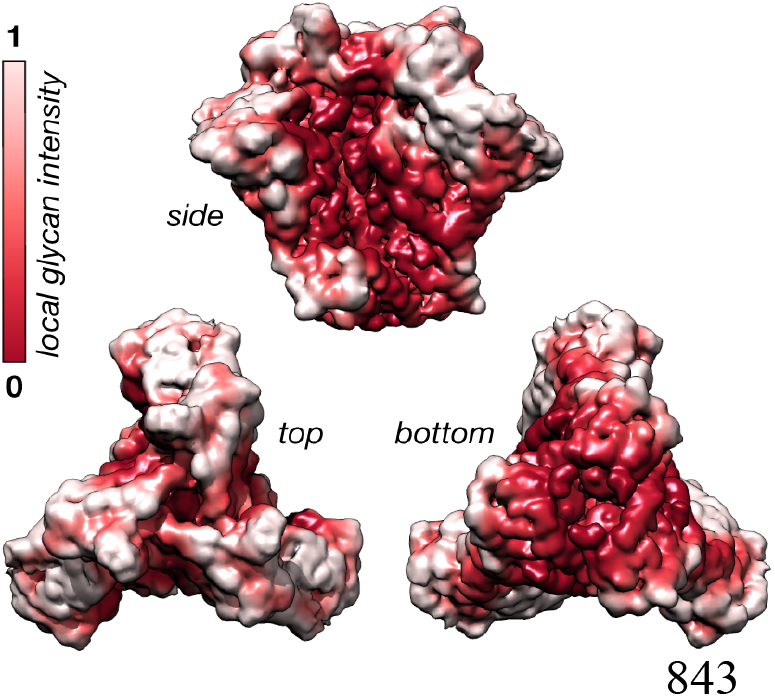
Quantifying the glycan shielding effect from cryo-EM maps. The normalized glycan shielding effect for BG505_CHO defined as the total glycan signal within a ~7Å radius spherical probe centered at each voxel. White surfaces are strongly shield by glycans.

Looking beyond the results of this paper, our integrated approach can be easily extended to other Env clones and to heavily glycosylated spike proteins from other viruses such as Influenza, Ebola, Lassa, and Coronaviruses, and represents a potentially powerful approach for studying the structure and dynamics of glycoproteins in general.

## Materials and Methods

BG505 SOSIP.664v3 was expressed and purified from HEK293F and HEK293S suspension cell culture in house while the CHO cell-line derived BG505 SOSIP.664 was provided to us by IAVI and expressed and purified as described in Dey, et al., 2017 (31). The monoclonal antibody RM20A3 was isolated from a BG505 SOSIP.664 immunized rhesus macaque(62) and the Fab was expressed and purified from HEK293F cells as described previously(63). The BG505 SOSIP.664 samples expressed in HEK293F and CHO cell lines were prepared for MS analysis as described previously(25), and the HEK293S sample was prepared with slight modifications on that protocol. All samples were prepared for cryo-EM analysis via the same protocol after being complexed with RM20A3 Fab. Imaging was performed on either an FEI Titan Krios or Talos Arctica (Thermo-Fisher) microscope. All non-custom cryo-EM data processing steps were performed using a combination RELION-2/3(64, 65) and CryoSparc v1(66). Model building and refinement was carried out with UCSF Chimera (67), Coot (68, 69), and Rosetta(70). All custom analysis of cryo-EM maps were performed in MatLab (2018b) {MATLAB and Image Processing Toolbox Release 2018b, The MathWorks, Inc., Natick, Massachusetts, United States.} 3-D variability analysis was performed in SPARX(27). The HT-AM was carried out by implementing the ALLOSMOD(29, 71) package of MODELLER(72, 73) in a streamlined pipeline. All graph theory and network-based analysis were performed using Python {Python Software Foundation, https://www.python.org/} and Matlab_R2018a packages. Simulated cryo-EM data generation and analysis was performed in RELION. All cryo-EM map and model visualization was performed with UCSF Chimera. A comprehensive description of all the methods can be found in the *SI Appendix*.

## Acknowledgements

Z.T.B would like to acknowledge Bill Anderson and Hannah Turner for assistance with microscopy, Charles Bowman and JC Ducom for computational support, Pawel Penczek, Francisco Asturias, Jack Johnson, and Gabe Lander for helpful discussions on 3-D variance analysis and cryo-EM data processing, Jesper Pallesen and JH Lee for early support with data processing, Jeffrey Copps for help with protein expression and purification, Lauren Holden for critical reading of the manuscript, and finally IAVI, Antu Dey, and John P. Moore for supplying demo run material and MS analysis. S.C and S.G. acknowledge Timothy Travers for proposing ALLOSMOD for this work, Nicolas Hengartner for helpful discussions on graph theory and Bette Korber and Kshitij Wagh for insights on glycosylations in HIV. S.C. and S.G. acknowledge the LANL High Performance Computing Division for providing computational facilities. We thank Robin Sung Kyu Park and Titus Jung for help with analysis of MS data for site specific glycosylation.

## Additional Information

### Author contributions

Conceptualization, Z.T.B, S.C., C.A.L, S.G. and A.B.W.; Methodology, Z.T.B, S.C., C.A.L, X.W., J.C.P., S.G. and A.B.W.; Protein expression and purification, Z.T.B., C.A.C, and J.L.T; Antibody isolation, M.J.vG; Mass-spectrometry experiments and data analysis, X.W, J.K.D, J.R.Y., and J.C.P.; Cryo-EM experiments, Z.T.B.; Modeling and Simulations, S.C.; Formal Analysis, Z.T.B. and S.C.; Visualization, Z.T.B. and S.C.; Manuscript Preparation, Z.T.B, S.C., C.A.L, J.C.P., S.G. and A.B.W.; Supervision, S.G. and A.B.W.

### Data Availability

All data needed to evaluate the conclusions in the paper are present in the paper and/or the Supplementary Information. Additional data related to this paper may be requested from the authors. The electron potential maps for BG505_293F, BG505_CHO, BG505_293S, BG505_EndoH2, and BG505_EndoH16 have been deposited in the Electron Microscopy Data Bank under accession codes 22108, 22109, 22110, 22111, and 22112 respectively, along with all corresponding half-maps, masks, and SPARX 3-D variability maps. Atomic models have been deposited in the Protein Data Bank under accession codes 6X9R, 6X9S, 6X9T, 6X9U, 6X9V.

