## Supplemental Appendix for "Visualization of the HIV-1 Env Glycan Shield Across Scales"

### Supplemental Information

#### An auto-thresholding method for cryo-EM maps based on topological connectivity

For visualization and interpretation, 3-D scalar field data such as cryo-EM maps are often displayed as an isosurface, which is a surface constructed form all the voxels of equal intensity, and the chosen intensity is often referred to as the *threshold*. The shape and topology of the isosurface can change dramatically with intensity and thus the choice of threshold will affect the interpretation of the data. In the case of cryo-EM maps, which consist of bright features in a noisy background, the best threshold is usually one that maximizes the visible signal while minimizing the amount of low intensity noise. It is also important that the resulting isosurface captures the topology of the molecule as accurately as possible. Therefore, for our scale-space analysis we implemented an auto-thresholding method based on *connected components*. Here, a connected component is any fully connected and isolated feature within an isosurface or binary volume. In the language of topology, the number of connected components defines the *0th Betti number*(1), where higher numbers correspond to more fragmented surfaces. When this number is calculated as a function of threshold (from high to low) the resulting curve is bimodal (Supplemental Figure 2A). Here, we demonstrate the method using the sharpened 3.1Å-resolution BG505_293F map. The first peak (left) corresponds to high-intensity signal while the second sharper peak (right) corresponds to low-intensity noise. We defined the *noise threshold* as the global minimum between these two peaks, which can be thought of as the maximally connected state before the appearance of low-intensity noise (red circle). This method differs from most auto-thresholding methods that operate on the distribution of pixel/voxel intensities, such as the popular Otsu's method(2). Supplemental Figure 2B shows the voxel intensity histogram for BG505_293F along with the thresholds determined by the topological method described here and Otsu's method (dashed vertical lines), while Supplemental Figure 2C shows the resulting isosurfaces. Both methods capture all the relevant signal at this scale, however, Otsu's method underestimate the threshold, including too much low intensity noise. When the map is smoothed with a Gaussian kernel first (SD = 2), both methods accurately exclude low intensity noise, but Otsu's method picks a slightly higher threshold, reducing the amount of low intensity signal that is visible and underestimating the volume of the map.

#### SPARX 3-D variability analysis and validation with the simulated cryo-EM data

Determining spatial variance in a cryo-EM map can be useful for identifying if there is sub-occupancy of bound ligands and for visualizing dynamic regions of macromolecules or macromolecular complexes. Here, we utilized 3-D variability analysis in the SPARX software package(3, 4). The method involves grouping projections with similar Euler angles, which were determined previously during 3-D refinement, and calculating a set of 2-D variance fields from these grouped projections (5). The 2-D variance fields are then used to reconstruct a 3-D volume with same algorithm used for standard 3-D reconstruction. Because this algorithm enforces positivity and the covariance term in the calculation can be positive or negative, the resulting map is not a true 3-D variance and is thus referred to as *3-D variability*. This can lead to some differences in the resulting map relative to a true 3-D variance. In addition, grouping 2-D projections with slightly different Euler angles will inevitably cause some 3-D rotational artifacts, although this can be reduced by selecting tighter groupings. However, the groups must also be large enough to minimize noise, so a balance has to be found between noise and rotational artifacts. As such, the number of particles in the dataset and the uniformity and completeness of the angular distribution will strongly influence the results. To help diagnose rotational artifacts, a 3-D average map is also calculated from the grouped projections. Here, the inclusion of the RM20A3 Fab resulted in a highly uniform and near-complete Euler angle distributions (Figure 1C), and the dataset was large enough (~60K particles) that the group size could be kept relatively large (50 projections/group) to minimize noise without causing significant rotational artifacts.

To determine how well the SPARX 3-D variability map captures the true 3-D variance in the glycan shield and diagnose any potential rotational artifacts, we calculated the true 3-D variance map for the simulated Man9 dataset. Supplemental Figure 7A-B shows a side-by-side comparison of the true per-voxel 3-D variance calculated from the simulated BG505_Man9 ensemble and the SPARX 3-D variability map for the same dataset. The variability map was calculated using the alignment parameters taken from refinement in RELION, and we used the same group size used for the experimental variability map (50 projections/group). It is clear that the two maps are nearly identical after being filtered to similar scales. The simulated Man9 dataset is of similar size and have a similarly uniform angular distribution as the experimental data, thus we concluded that the experimental variability maps should also be free from major distortions.

##### Determining convergence of the ALLOSMOD based high-throughput modeling pipeline

To determine if the HT-AM ensembles had converged, we calculated the root mean-squared fluctuation (rmsf) for each glycan across all 1000 models after aligning the protein scaffold (Figure 3E), then compared it to the average of randomly selected equally sized subsets (Supplemental Figure 5D). We see that the mean rmsf values between the subsets are nearly identical and the standard deviations from the means are very small, indicating convergence. A similar trend can be seen for the per-glycan sampled volumes (Supplemental Figure 5E).

##### Estimating the glycan shielding effect from cryo-EM maps

The cryo-EM maps presented here approximate the time-averaged structure of a single fluctuating Env molecule, and thus provide a direct experimental mapping of the shielding effect created by the dynamic glycans. To quantify this more precisely we developed a simple "rolling sphere" method that calculates the total glycan signal within a spherical probe centered at each voxel in the map. To isolate the glycan signal from the protein we performed signal subtraction with the BG505_EndoH16 map (Supplemental Figure 18A). Because BG505_EndoH16 still contained some residual glycan signal from incomplete digestion, this was first removed via masking. Finally, a soft mask in the shape of the RM20A3 Fabs was used to remove any Fab signal remaining after subtraction. A spherical probe was then rastered across this glycan-only difference map and the total intensity within the sphere was recorded at every voxel (Supplemental Figure 18A). This cumulative signal intensity can be seen as representing the relative shielding at every point in the map, with higher local glycan intensity representing stronger shielding. We chose a ~7Å radius probe to imitate the extended loops commonly employed by neutralizing antibodies to penetrate the inner layers of the glycan shield (6, 7). The final step involves determining the proper normalization, however, there is no clear way to do this objectively. Therefore, we adjusted the scale such that surfaces which are still exposed when the 2 SD Gaussian filtered glycan-only difference map is visualized at its noise threshold are set to 0 (red), and the surfaces occluded by a single glycan on average are set to 1 (white - Supplemental Figure 18B). In Figure 11 and Supplemental Figure 18B we show the results of this analysis for BG505_CHO projected as a linear color map onto the surface of the modified BG505_EndoH16 map used for signal subtraction (the RM20A3 Fabs were removed by masking to expose the base of the SOSIP trimer). White surfaces are strongly masked while red surfaces are weakly masked, highlighting potential sites of vulnerability to neutralizing antibodies. Because this method calculates local glycan signal and not surface accessibility, regions that are occluded by protein but not glycans, such as the interior of the structure and the protomer-protomer interfaces, are also colored red despite being obviously inaccessible to antibodies.

#### Materials and Methods

##### BG505 SOSIP.664v3 expression and purification in HEK293F, 293S, and CHO cells

BG505 SOSIP.664v3 was expressed and purified from 1L HEK293F and HEK293S suspension cell cultures by the following protocol. First, cell cultures were expanded then transiently co-transfected with Env and Furin plasmid DNA in a 1:3 (w/w) ratio with PEI. Expression was carried out for 6 days followed by supernatant harvesting by centrifugation and protein purification via PGT121 antibody affinity and size-exclusion chromatography in 1X TBS buffer. Fractions corresponding to stable trimer were pooled and concentrated. Demo run BG505 SOSIP.664v3 was provided to us by IAVI and was expressed and purified from a stable CHO cell-line as described in Dey, et al.(8)

##### RM20A3 IgG isolation and Fab purification

The monoclonal antibody RM20A3 was isolated from a BG505 SOSIP.664 immunized rhesus macaque (rh2011) at week 53, which was 1 week after the 6$^{th}$ immunization(9), using single cell FACS sorting and antibody cloning as previously described(10). A full description of the antibody will be presented in elsewhere. The RM20A3 Fab was expressed and purified from HEK293F cells as described previously(11).

##### Site-specific mass spectrometry

The BG505 SOSIP.664 samples expressed in HEK293F and CHO cell lines were prepared for MS analysis as previously described(12), and the HEK293S sample was prepared with slight modifications on that protocol. In brief, since glycans from HEK293S cells are all oligomannose (Man5-Man9) BG505 SOSIP.664 samples peptide mixtures were deglycosylated only with endoglycosidase PNGase F in 100 mM ammonium bicarbonate prepared with O18-water.
The samples were analyzed on an Q Exactive HF-X mass spectrometer (Thermo). Samples were injected directly onto a 25 cm, 100 μm ID column packed with BEH 1.7 μm C18 resin (Waters). Samples were separated at a flow rate of 300 nL/min on a nLC 1200 (Thermo). Solutions A and B were 0.1% formic acid in 5% and 80% acetonitrile, respectively. A gradient of 1–25% B over 160 min, an increase to 40% B over 40 min, an increase to 90% B over another 10 min and held at 90% B for 30 min was used for a 240 min total run time. Column was re-equilibrated with solution A prior to the injection of sample. Peptides were eluted directly from the tip of the column and nanosprayed directly into the mass spectrometer by application of 2.8 kV voltage at the back of the column. The HFX was operated in a data dependent mode. Full MS1 scans were collected in the Orbitrap at 120k resolution. The ten most abundant ions per scan were selected for HCD MS/MS at 25NCE. Dynamic exclusion was enabled with exclusion duration of 10 s and singly charged ions were excluded.

The MS data were processed as previously(12). The data were searched against the proteome database and quantified using peak area in Integrated Proteomics Pipeline-IP2. For samples produced in HEK293F and CHO cells, glycosites (N-X-T/S) with N + 203 were identified as sites with high-mannose glycans removed by the initial Endo H treatment (high-mannose), the glycosites with N + 3 were identified as sites whose glycans were complex type glycans removed by PNGase F, and glycosites with N+0 were identified as sites that had no glycan prior to endoglycosidase treatments. Since samples produced in HEK293S are only high-mannose, and were treated only with PNGase F, sites with N+3 were identified as sites with high-mannose.

**Endoglycosidase H digestion of BG505 SOSIP.664 for cryo-EM experiments**

1.4 mg of purified BG505 SOSIP.664 from HEK293S cells in complex with RM20A3 was mixed with 80,000 units of endoglycosidase H (Endo H; New England Biolabs) in non-denaturing reaction buffer to a final volume of 1ml and incubated at 37$^{o}$C for 2 hrs and 16 hrs (two separate reactions from the same master mix). To quench the reaction and purify the sample for cryo-EM experiments, it was run over an SEC column and fractions were pulled and concentrated. When the sample was used for MS or SDS PAGE, the reaction was heat quenched at 100°C for 10min.

**Cryo-EM sample preparation for BG505 SOSIP.664 in complex with RM20A3**

Purified BG505 SOSIP.664v3 from either HEK293F, HEK293S, or CHO cells was incubated overnight at 4°C in a 1:6 molar ratio of SOSIP trimer to purified RM20A3 Fab. The complex was purified from the remaining free Fab by SEC, then concentrated to ~6mg/ml in 1X TBS buffer. Purified complex was then mixed with n-dodecyl β-D-maltoside (DDM; Anatrace) to a final concentration of 0.06 mM immediately prior to vitrification. 3ul of this solution was then applied to plasma cleaned Cflat 1.2x1.3 holey carbon grids and plunge frozen at 4°C and 100% chamber humidity with an FEI Vitrobot MarkIV (Thermo-Fisher) after blotting for 7 s. Grids were transferred in liquid nitrogen to an autoloader and into one of the two microscopes for imaging. The Endo H digestion samples were prepared in the same way after purification from the reaction mixture.

**Cryo-EM data collection**

Imaging was performed on either a FEI Titan Krios or Talos Arctica (Thermo-Fisher), operating at 300KeV and 200KeV respectively. The microscopes were taken through a standard coma-free alignment protocol before each imaging session. Movie micrographs were collected on a Gatan K2 Summit direct electron detector (Gatan) operating in counting mode. Imaging was adjusted to achieve the same total dose for both microscopes of ~50 e-/Å$^2$. Movies were collected at a frame rate of 1/250 ms. The final calibrated pixel sizes were 1.03 Å for the Krios and 1.15 Å for the Arctica. During data collection, frames were aligned and dose weighted in real-time with MotionCor2 (13) and CTF fits were performed using Gctf(14) to monitor image quality.

**Cryo-EM data processing**

All non-custom cryo-EM data processing steps, which included particle picking, 2-D and 3-D classification, refinement, per-particle CTF refinement, and postprocessing were performed using a combination RELION-2/3(15, 16) and CryoSparc v1(17). Micrograph sorting and carbon masking were conducted with EMHP(18) when necessary. The following general processing protocol was used for all datasets. After global CTF estimation, all micrographs with resolution estimates greater than 6Å were excluded from the dataset. Particle picking was then performed using the Gaussian disk template in RELION-2. Extracted particles were then transferred to CryoSparc 1 servers for initial 2-D and 3-D processing. The initial processing usually involved 2 rounds of 2-D classification and subset selection followed by 1 round of 3-D ab-initio refinement into 4 classes. Clean classes were then pooled and refined using the class average as a template. All refinements were performed with C3 symmetry unless stated otherwise. At this stage, refinement meta-data was downloaded from CryoSparc and re-formatted for processing in RELION. Following refinement in RELION, per-particle CTF and beam-tilt refinement were performed, followed by another round of refinement and 1 or more rounds of 3-D classification using a tight mask and local angular searches only. For the Endo H treated samples, subtractive classification in RELION-3 was also performed at this stage to focus classification to the heterogenous apex. Final sorted particles were then refined using C3 symmetry and sharpened with a data-derived B-factor in RELION using the mask from refinement. Map segmentations were performed with Segger(19) in UCSFChimera(20). Local resolution plots were calculated in RELION using the method presented in Kucukelbir et al., 2014 (21).

**Model building and refinement**

Model building and refinement was initially performed on the sharpened 3.1Å-resolution BG505_293F map using the following protocol: First, models of the complete gp120, gp41, and the RM20A3 heavy and light chain variable domains, were generated with SWISS-MODEL(22) using PDBID:5ACO as the Env template. These models were then docked into the sharpened map and saved as a single PDB file using UCSF Chimera(20). This model was then refined with C3 symmetry in Rosetta(23), asking for 300 models. Each model was scored with MolProbity (24) and EMRinger (25), and the one with the best overall score was selected. N-linked glycans were then modeled into the map using Coot(26) and the glycosylated model was re-refined in Rosetta using the newly introduced glycan force fields (27). After refinement, coordinates outside of clear density were deleted and manual adjustments to the structure were made with Coot where necessary, followed by another round of refinement in Rosetta. This final BG505_293F model then served as the input for a single round of Rosetta refinement into the BG505_CHO, BG505_293S, BG505_EndoH2, and BG505_EndoH16 maps. Full model C$\alpha$ RMSDs were calculated in PyMol(28) {The PyMOL Molecular Graphics System, Version 1.2r3pre, Schrödinger, LLC.}.

**Auto-thresholding and scale-space analysis**

For all quantitative comparisons between the cryo-EM maps, either experimental or simulated, which included scale-space analysis, local intensity analysis, and difference mapping, all maps were first aligned and resampled onto the same grid using the UCSFChimera function *vop resample*, then intensity equalized with the RELION-3 function *adjust_power*. Because the datasets were collected on two microscopes operating at different magnifications, all maps were resampled to the larger voxel size of 1.15Å and adjusted to match the power spectrum of lowest resolution reconstruction (BG505_EndoH16 - 3.5Å). Resampled and intensity equalized cryo-EM maps were loaded into MATLAB (2018b) {MATLAB and Image Processing Toolbox Release 2018b, The MathWorks, Inc., Natick, Massachusetts, United States.} using the *ReadMRC.m* script obtained from the following GitHub: https://github.com/Sarofi/PythonImages/blob/master/Matlab/FredSigworth/EMIODist/ReadMRC.m. All subsequent analysis was performed in MATLAB and the MATLAB ImageProcessing Toolbox. To determine the noise threshold, the maps were binarized at 100 evenly spaced intensities ranging from the maximum to minimum map intensity with the MATLAB function *imbinarize*, and the number of connected components of each binary volume was calculated using the MATLAB function *bwconncomp*. The noise threshold was then defined as the global minimum between the high and low intensity peaks (Supplemental Figure 3A,D). If the function plateaued at the minimum or the minimum was degenerate, the lowest intensity point in the plateau or lowest intensity minimum were selected (Supplemental Figure 3D). The volume of the map at the noise threshold was then calculated as the sum of the volumes of all the connected components measured in Å$^3$. The noise threshold and total volume were then calculated as a function of Gaussian filter width (measured in standard deviation - SD) to create the plots shown in Figure 2B and Supplemental Figure 3G-J. The analysis was also repeated for maps that had not been intensity equalized for comparison (Supplemental Figure 3H). To isolate the contribution from the glycan signal from that of the poorly resolved constant domains of the Fabs, the analysis was repeated after the removing the Fab density by masking (Supplemental Figure 3I), which showed reduced volume at low resolutions but no apparent differences in the shape of the curves nor the resolution scale of the inflection points and plateaus. For our implementation of Otsu's method, we used the MATLAB function *graythresh*.

**SPARX 3-D variability analysis**

3-D variability analysis was carried in the SPARX software package(3). Refinement *data.star* files from the final refinement in RELION (before per-particle CTF refinement) for each dataset were first converted into SPARX format with the function *sxrelion2sparx*, then 3-D variability maps were calculated with the function *sx3dvariability* using C3 symmetry. Prior to calculating 3-D variability, projections were internally down sampled 2x and lowpass filtered to twice the binned Nyquist frequency to reduce noise in the resulting variability map. The resulting raw 3-D variability maps were too noisy to interpret, so they were smoothed with a 2.5 SD Gaussian kernel. To calculate the true per-voxel 3-D variance and 3-D average from the BG505_Man9 simulated cryo-EM dataset, simulated volumes were generated for each pre-aligned model using the EMAN2 function *e2pdb2mrc* then loaded into . For each voxel in the maps, the variance and mean intensity were calculated and saved as mrc volumes with the MATLAB function *WriteMRC* obtained from the following GitHub: https://github.com/Sarofi/PythonImages/blob/master/Matlab/FredSigworth/EMIODist/WriteMRC.m.

**Cryo-EM difference mapping**

Difference maps were calculated by subtracting one map from another in the order mentioned in the text using the UCSFChimera(20) function *vop subtract* followed by smoothing with a Gaussian kernel for visualization.

**Measuring local map intensity around glycan BMA residues**

To measure the relative signal intensity around each individual glycan in the cryo-EM maps, first the maps were filtered with a 1.5 SD Gaussian kernel and saved in UCSFChimera. Filtered maps there then opened in Coot along with the corresponding refined atomic models and a full idealized N-linked glycan stem (first three glycan residues) was added at every site that could be confidently identified, if not already present, and adjusted to fit the density with the *refine tree* function(26). To determine how well the dynamics at each glycan residue approximated the average dynamics of the entire glycan, we calculated the mean deviation from the full glycan average rmsf for each of the 11 Man9 residues (Supplemental Figure 8A). In addition, we calculated the Pearson correlation coefficient between the per-glycan average rmsf when using the full glycan and when using each residue alone (Supplemental Figure 8B). Of the glycan residues we could confidently build into the cryo-EM maps, the BMA residue most closely approximated the full glycan average. Therefore the average X,Y, and Y coordinates of each BMA residue were then used to localize each glycan within the volume, and the average voxel intensity within a spherical probe around each BMA residue was calculated for a range of probe sizes along with the Pearson correlation coefficient between the normalized inverse rmsf and normalized mean intensities (Figure 5B). The same model was used to analyze the BG505_293F and BG505_CHO maps, while a minor adjustment to the N611 glycan was made for the BG505_293S map. The BG505_293S model was also used to analyze the two Endo H digestion intermediate maps. This same procedure was used to analyze the BG505_Man9, Man9HO, and Man5 simulated cryo-EM maps. To quantify the percent occupancy at each site after Endo H digestion we assumed a linear relationship between signal intensity and occupancy, with any intensity <= 0 being considered fully digested, and the initial occupancies were determined by the MS data for the BG505_293S sample. To calculate the Endo H protection score we added the percent occupancy at each site from the EndoH2 and EndoH16 maps and normalized the results between 0 and 1.

**Glycan crowding analysis**

We defined a simple crowding score as the number of glycans within a spherical probe around each glycan BMA residue. To localize each glycan, we calculated the average location of the BMA oxygen atom across all 1000 models in the Man9 ensemble. Then, the number of glycans within a spherical probe around each point was recorded as a function of radius (0Å-135Å, where 135Å was the maximum pairwise distance interaction observed in the ensemble) along with the Pearson correlation between the normalized crowding score and either the normalized inverse rmsf, normalize local map intensity, normalized percent high-mannose, normalized Endo H protection, or normalized eigencentrality (Supplemental Figure 6).

**ALLOSMOD-based high-throughput atomistic modeling (HT-AM) of fully glycosylated Env**

We generated a robust computational ensemble of atomistic models of BG505 SOSIP.664 by implementing the ALLOSMOD(29, 30) package of MODELLER(31, 32) in a streamlined pipeline. The BG505 SOSIP.664 protein scaffold was homology modeled and the missing residues in the hypervariable and dynamic V2 and V4 loops (residues 186-189 and residues 400-410) were modeled *ab initio*. All the disulfide bonds were maintained as additional restraints. 100 models were generated, and the best 10 were selected (Figure 3A) based on MODELLER optimization scores and stereochemistry scores as determined by PROCHECK(33). Due to the long stretches of missing residues in loops V2 and V4 (residues 185A-185I and 400-410) these loops were modeled template-free and the 10 structures sample different loop conformations, accounting for their structural variability. For each of the 10 selected protein structures, glycans were initially added at the known glycosylation sites based on ideal geometries as dictated by CHARMM36 (34, 35) force field internal coordinates. For simplicity, we chose to apply uniform glycosylation of mannose-9 because the MS data suggests this is the most common glycan type on BG505 SOSIP.664(36). This was followed by a 1Å random deviation added to the overall atomic coordinates. This template-free glycan modeling method optimizes an energy function given by a combination of spatial restraints and CHARMM36 glycan forcefield terms, to enforce proper stereochemistry. The generated structures were further relaxed (Figure 3B) with several steps of conjugate gradient minimization followed by simulated annealing (SA) as described by Guttman et al. (29).

The glycans were kept completely flexible during the refinement steps, and the protein backbone was harmonically restrained to the template, to prevent local unfolding during the SA steps. 100 fully glycosylated structures were modeled from each of the 10 selected protein models, resulting in the final ensemble containing 1000 different poses.

**Rmsf and sampling volume calculations from HT-AM ensembles**

For both the native and all-man9 glycosylated models, the root mean square fluctuations (rmsf) of each glycan *n* was calculated as an average over all its heavy atoms, by the following equation:

$$RMSF_n = \frac{1}{K}\sum_{k=1}^{K}\sqrt{\frac{1}{M}\sum_{m=1}^{M}|\vec{r}_{mnk} - \langle\vec{r}_{nk}\rangle|^2}$$

where $\vec{r}_{mnk}$ is the atomic position of heavy atom *k* of glycan *n* in snapshot *m*, $\langle\vec{r}_{nk}\rangle = (\frac{1}{M})\sum_{m=1}^{M}\vec{r}_{mnk}$ is the average atomic position of heavy atom *k* in glycan *n*. K is the total number of heavy atoms in the glycan (127 for Man9). The ensemble for each model contains 1000 snapshots, making *M* = 1000 snapshots for each of the two models. The standard deviations (s.d.) were obtained by dividing the 1000 snapshots into 4 sets of M=250, and calculating the square root of the variations from the four sets of rmsf values. The single-model average rmsf was similarly calculated, where M reduces to 100. The value obtained for each of the 10 models were averaged to get the final per-model rmsf value. For calculating the sampled volumes, volumetric surface maps were created for each individual glycan using the volmap plugin of VMD(37) (http://www.ks.uiuc.edu/Research/vmd/), including all 1000 models, with 1Å grid size. The enclosed volume of these generated surface models was calculated using UCSF CHIMERA (20) (http://www.rbvi.ucsf.edu/chimera).

**Graph theoretic analysis of ALLOSMOD ensembles**

The inter-glycan overlap is calculated as the total fraction of heavy atoms from the two glycans that come within 5Å of each other. A single mannose-9 glycan has 127 heavy atoms. Since our ensemble is composed of 1000 possible structures, there are effectively 127,000 heavy atoms per ensemble of mannose-9 at one position. The fraction of the total number of heavy atoms from two neighboring ensembles that come within contact distance defines the overlap fraction. An overlap greater than or equal to 50% of heavy atoms from two neighboring mannose-9 glycans is assigned as 1. This overlap matrix is used to define the adjacency matrix for our network analysis. Each glycan functions as a node of the graph, and two nodes are connected by an edge if there is at least 5% overlap as per our overlap definition given above. The edge length is inversely proportional to the overlap value, i.e., the larger the overlap, the closer two nodes (glycans) are in the graph. Only those glycans from the neighboring protomers are considered, that have an inter-protomer edge. All graph theory and network-based analysis were performed using Python {Python Software Foundation, https://www.python.org/} and MATLAB_R2018a packages.

**Simulated cryo-EM data creation and processing from HT-AM ensembles**

First, each model generated from the HT-AM pipeline was converted into *mrc* volume with the same box and pixel sizes as the experimental data (in this case we used the lower magnification from the Talos Arctica of 1.15Å/pixel) using the EMAN2(38) function e2*pdb2mrc*, with the max resolution set as twice the voxel size. Next, each volume was projected at 100 uniformly distributed angles with 100 SD white noise added using the RELION function *relion_project*. All 1000 *particles.star* files were then combined (100K total projections) and refinement was initiated using the BG505_293F map with Fabs removed as an initial model. In all cases the resolution converged to around the Nyquist frequency (2.3Å), and the final maps were sharpened using a data-derived B-factor and soft mask in RELION. SPARX 3-D variability maps were calculated in the same manner as the experimental data however without including CTF information. Because SPARX must re-extracts particles for variability analysis, the center of each projection image was given in place of the micrograph coordinates for each particle by adding *_rlnCoordinateX* and *Y* columns to the combined star file. True per-voxel average and variance maps were calculated by taking the mean and variance of each voxel across all 1000 *mrc* volumes used for synthetic data projections.

**Rolling sphere method for quantifying the glycan shielding effect from cryo-EM maps**

Resampled and intensity equalized cryo-EM maps were loaded into MATLAB and the glycan signal was isolated via difference mapping using a modified BG505_EndoH16 map with the residual glycan density removed. The difference map was then multiplied by a soft mask in the shape of the RM20A3 Fabs to remove any remaining Fab density. Next, a spherical probe with a radius of 6 voxels (6.9Å) was rastered across the difference map and the total intensity within the spherical probe centered at each voxel was recorded. The results were then saved as an mrc volume along with the modified BG505_EndoH16 map used for signal subtraction. These maps were then loaded into UCSFChimera and the local glycan intensity was used to color the surface of the modified BG505_EndoH16 map with a linear white/red gradient colormap.

#### Supplemental Figures

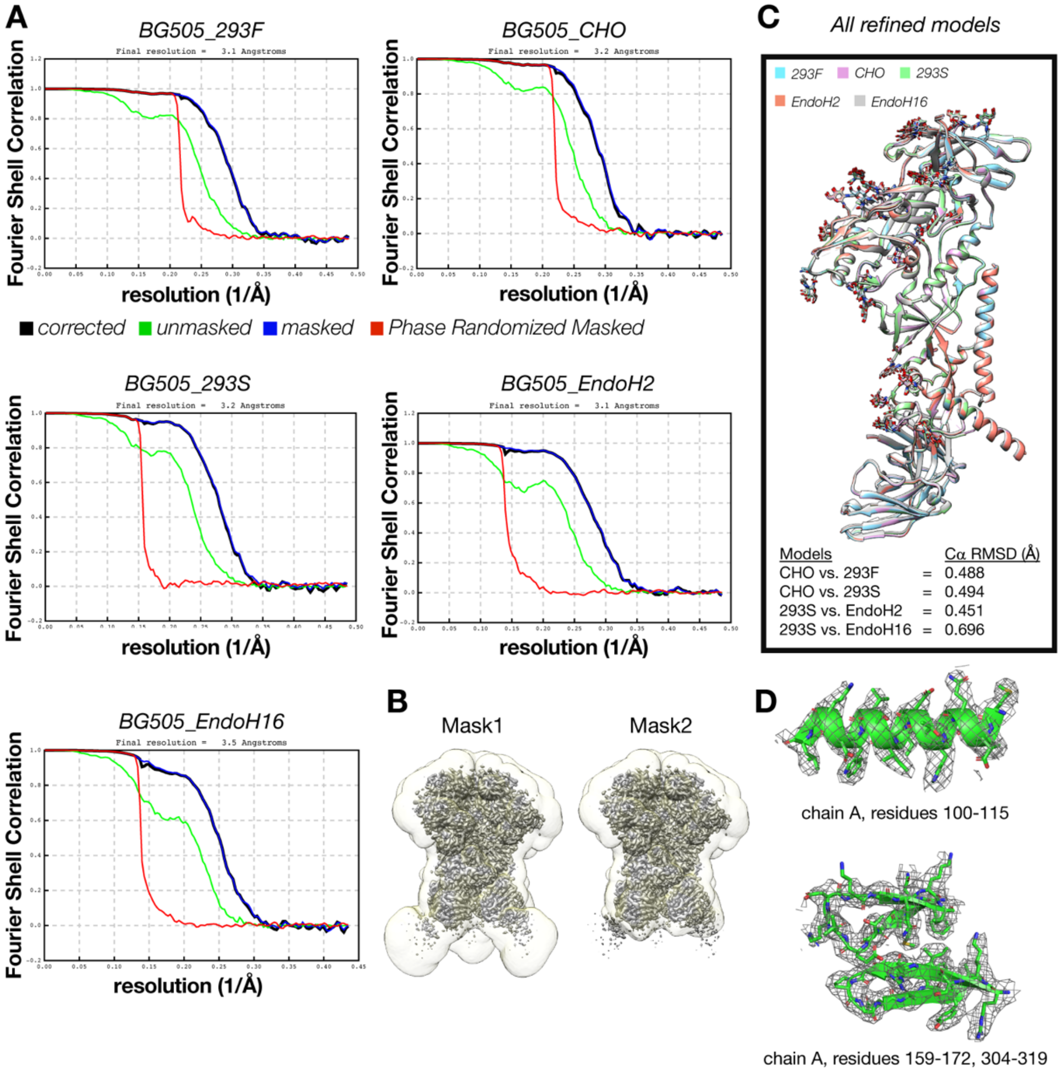


**Supplemental Figure 1 | Cryo-EM resolution plots and atomic models of BG505 SOSIP.664 in complex with RM20A3 Fab. (A)** Fourier Shell Correlation (FSC) plots for all five experimentally determined cryo-EM maps. **(B)** Soft masked used during refinement, classification, and sharpening. The disordered constant domains of the Fabs were masked out during later stages of refinement with Mask2. **(C)** All five refined atomic models superimposed along with alpha carbon root mean squared deviations (RMSD) between the listed pairs of models. **(D)** Two representative cryo-EM map density shots with underlying refined atomic models from the BG505_293F map.

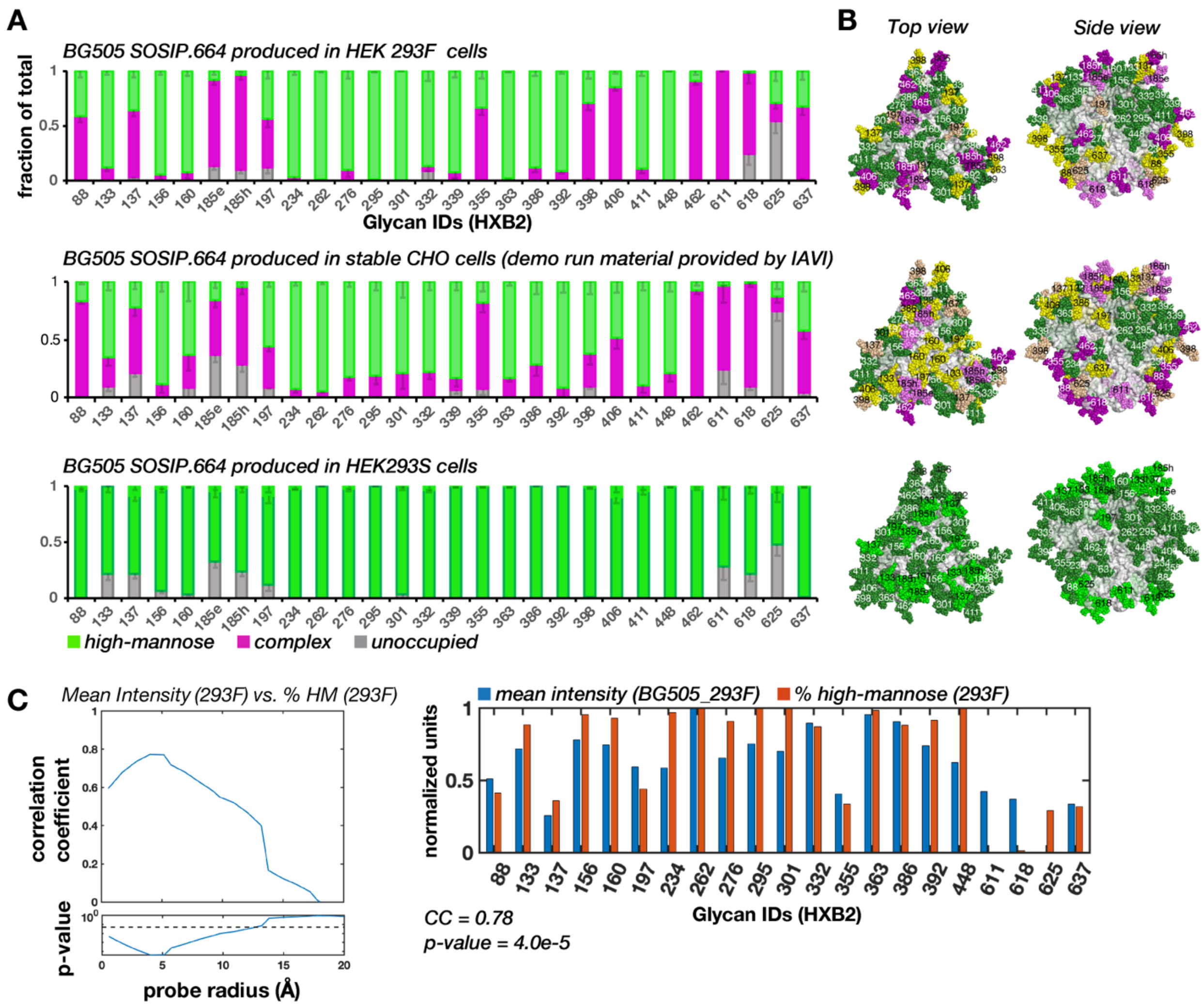


**Supplemental Figure 2 | Site-specific MS analysis of BG505 SOSIP.664 from 3 different cell lines. (A)** Site-specific MS profiles for BG505 SOSIP.664 expressed in 3 different cell types; HEK293F, CHO, and HEK293S. MS data shows the percentage total complex (magenta), high-mannose (green), and un-occupied (grey) at each site. Small green bars in the 293S data sample with green borders represent false-positive +203 mass signatures (see Methods). **(B)** Molecular surface representation of BG505 SOSIP.664 with an idealized Man5 glycan at each location colored according to composition. The color scale is as follows: If unoccupied >0.1, set lighter color, then recalculate the ratio of complex and high-mannose. If high-mannose >0.75 color green, if complex >0.75 color violet, if mixed color yellow. **(C)** Pearson correlation coefficient and p-value between the normalized mean local intensity around BMA residues and normalized percent high-mannose content at each site as a function of probe radius around BMA residues for the BG505_293F sample. The bar plot shows the site-specific intensity and percent high-mannose content at each glycan we could identify in the map using the probe radius with the peak correlation.

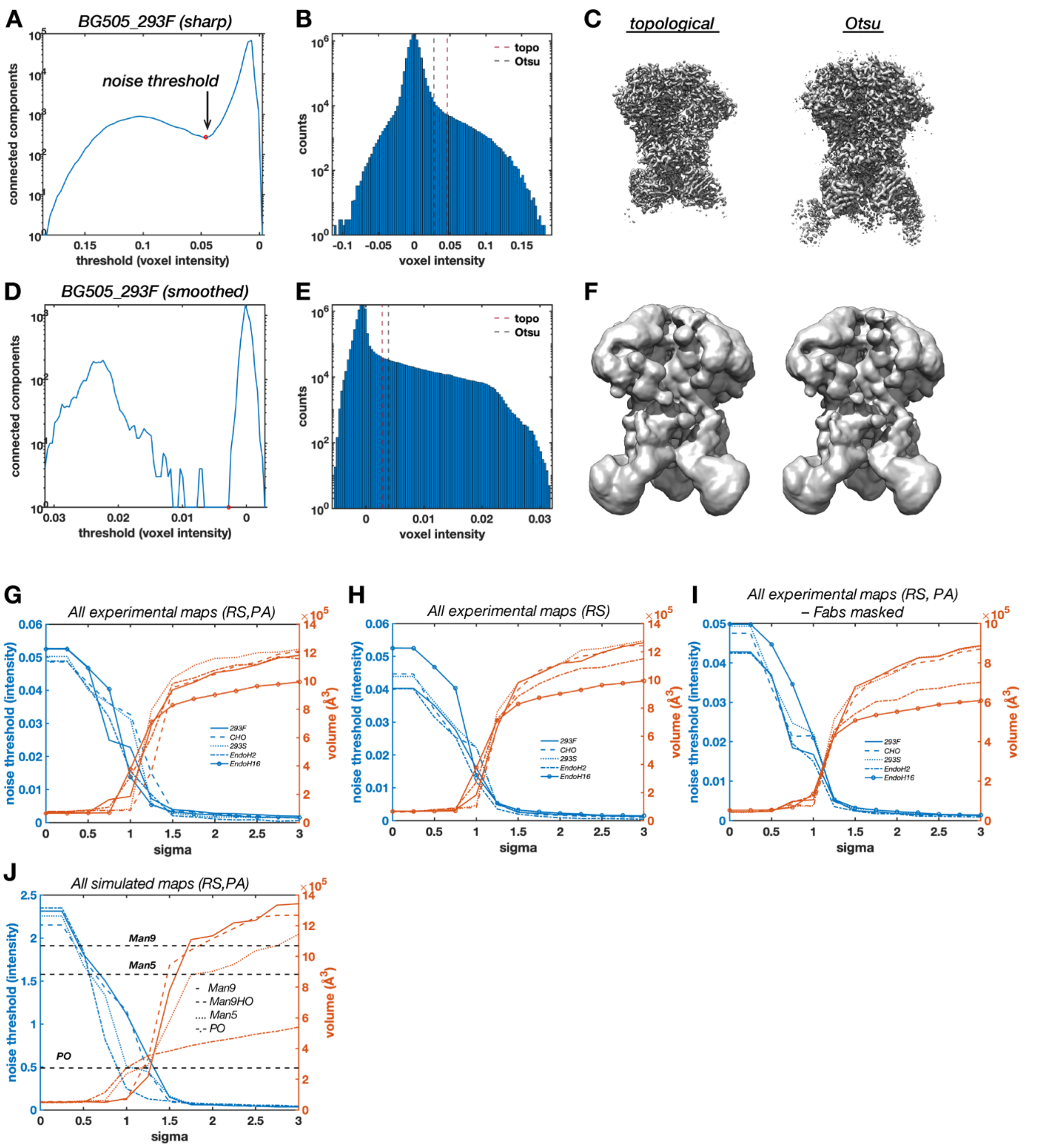


**Supplemental Figure 3| Auto-thresholding and extended scale-space analysis. (A)** The number of connected components as a function of threshold for BG505_293F. The red circle marks the noise threshold determined by finding the minimum between the high and low intensity peaks. **(B)** Voxel intensity histogram for BG505_293F showing the thresholds determined by the topological method described here and Otsu's method, along with the corresponding isosurfaces **(C)**. **(D-E)** Same as in panels A-C but after filtering the map with a 2 SD Gaussian kernel.**(G-H)** Scale-space analysis plots showing noise threshold and map volume at the noise threshold as a function of Gaussian filter width (measured in SD) for all experimental cryo-EM maps after being resampled to the same grid

(RS) and intensity equalized with the RELION function *adjust_power* (PA). **(I)** Same as in panel H but after masking out the RM20A3 Fabs. **(J)** Same as in panel G but for all simulated cryo-EM maps.

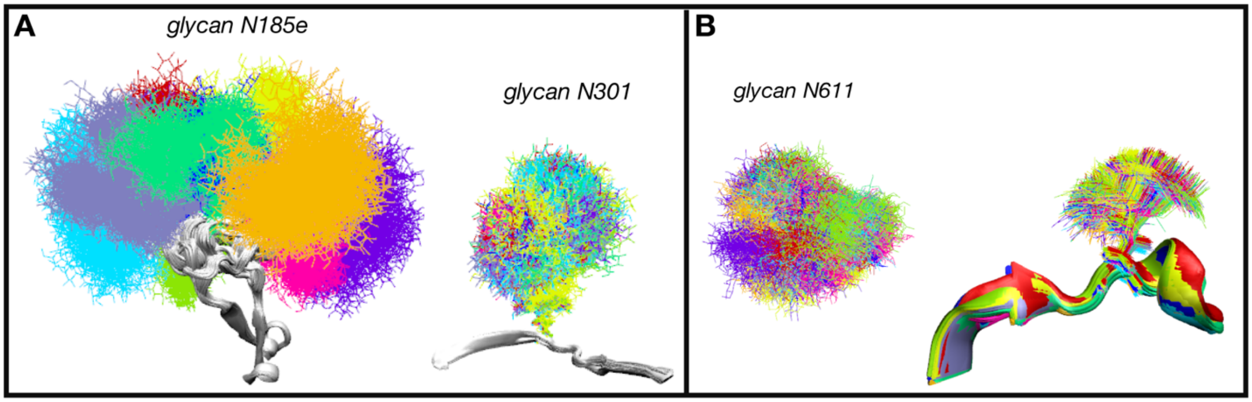


**Supplemental Figure 4 | Visualization of sampling volume at different glycosylation sites from the HT-AM Man9 ensemble. (A)** Comparative sizes of glycan volumes at positions 185e (high rmsf) and 301 (low rmsf). The sampling space of the underlying protein backbone is given in grey. Different colors are given for glycans from the 10 different starting protein models. **(B)** Top view of the different glycan models at residue 611 (left). Protein chain, Asn side chain, and first NAG ring for the different glycan

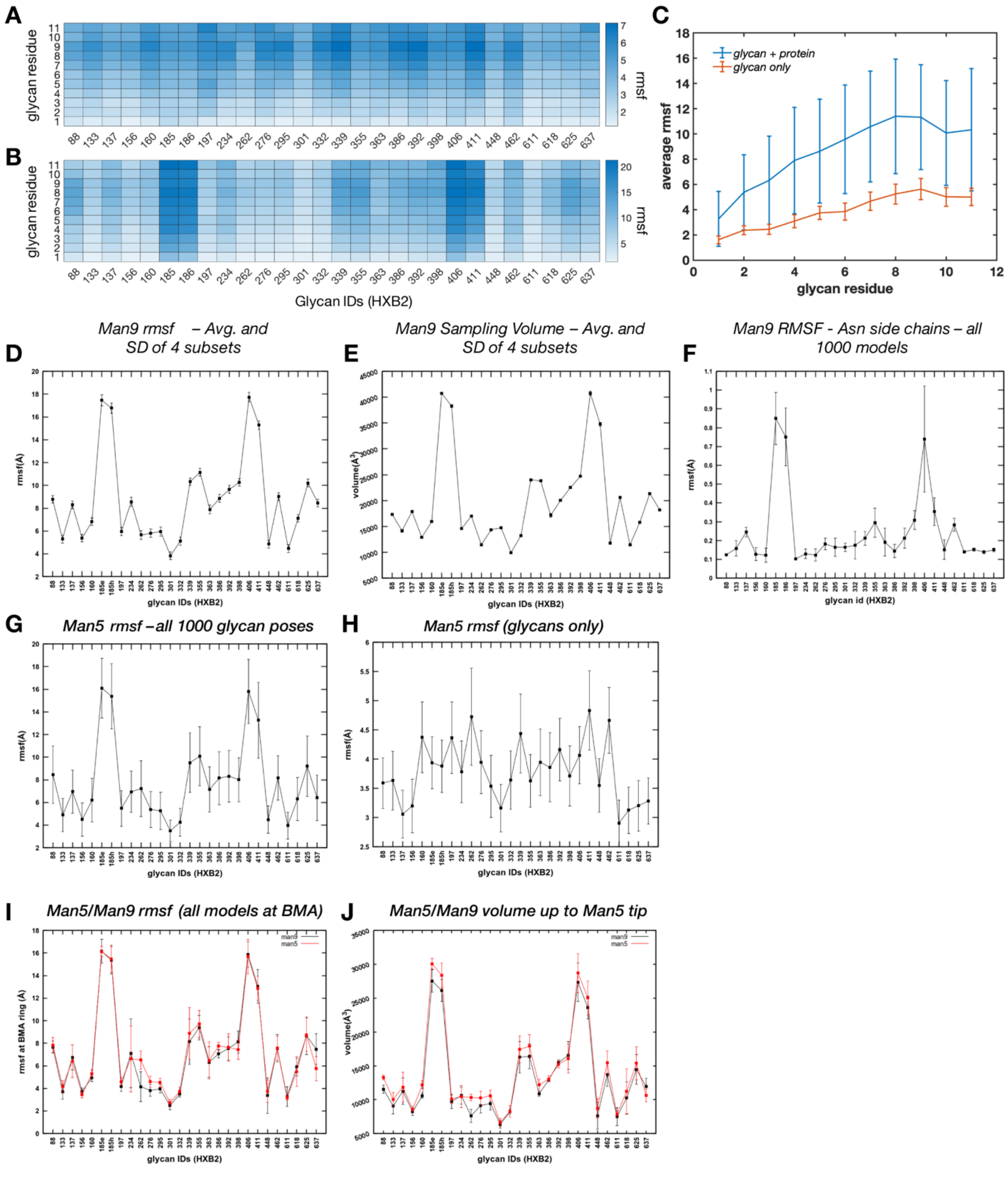


**Supplemental Figure 5 | End extended rmsf and volume sampling analysis for Man9 and Man5 HT-AM ensembles.** (A) Per-glycan residue all-atom mean rmsf for a single protein model (averaged over 10 sets of 100 glycan poses each) and **(B)** for all 1000 models. **(C)** Average rmsf for each individual glycan residue for the data shown in panels A and B. **(D)** Full glycan average rmsf and **(E)** sampled volume along with SD calculated over 4 sets of 250 poses each for the Man9 ensemble. **(F)** Average rmsf and SD at the glycosylated ASN side chains for the Man9 ensemble. **(G-H)** rmsf plots for Man5 ensemble as in Figure 3E-F. **(I)** Comparison of Man5 with Man9 all atom mean rmsf at the BMA residue. **(J)** Comparison of Man5 with Man9 sampled volume up to the 5$^{th}$ mannose residue. models at residue 611 (right). The Asn residue is located at the base of a helix. Colors depict the different starting protein models.

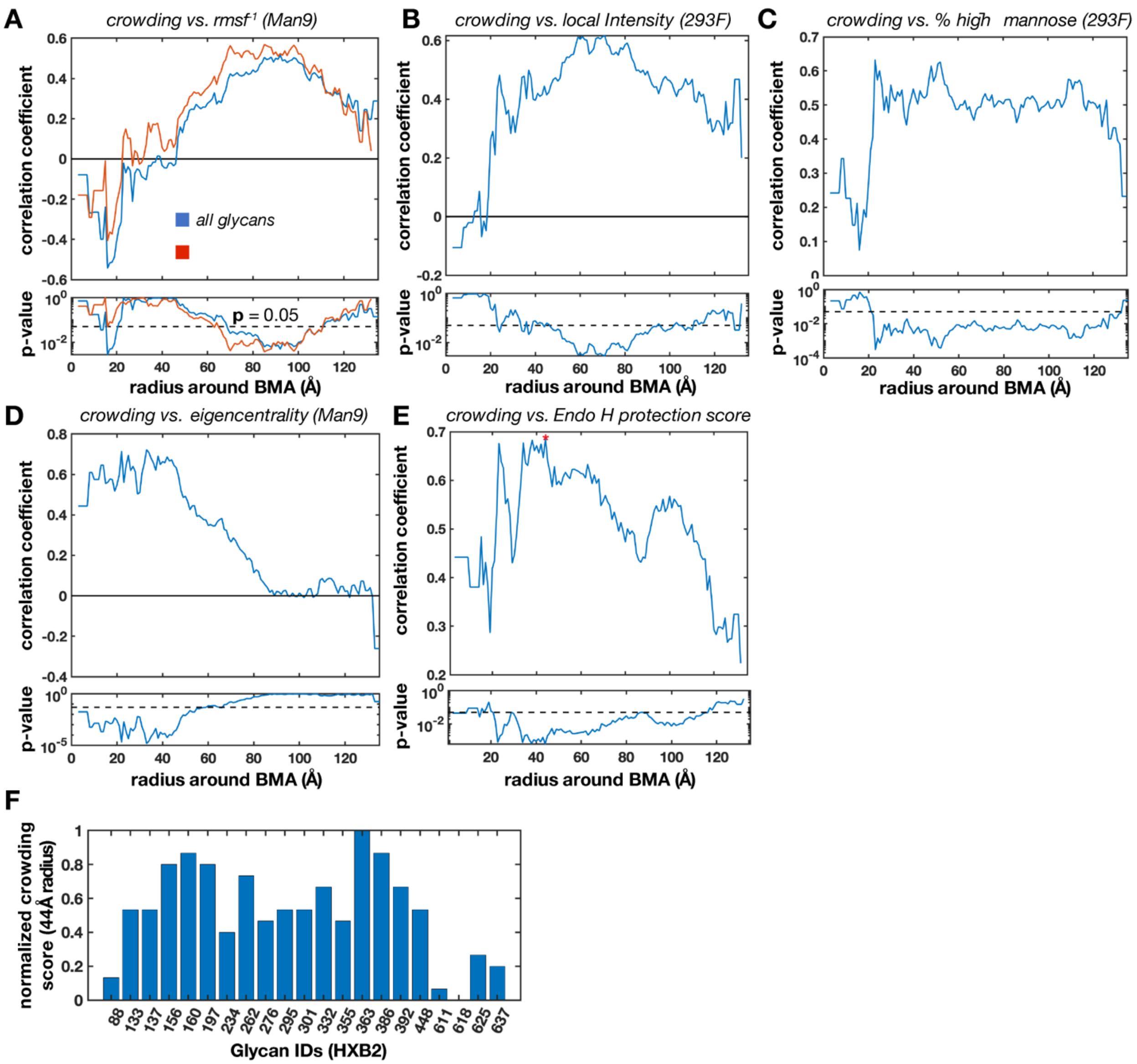


**Supplemental Figure 6 | Glycan crowding analysis. (A)** Pearson correlation coefficient and *p*-value between the number of glycans within a spherical radius around each BMA center of mass and inverse rmsf of the Man9 ensemble calculated considering every glycan (blue) and when excluding the highly dynamic V2 and V4 loop glycans (red) as a function of probe radius. **(B)** Same as in panel A except showing the correlation between crowding and the mean intensity at each glycan from the BG505_293F cryo-EM map (only those glycans we could identify), **(C)** the percentage of high-mannose type glycans in the 293F sample, **(D)** network eigencentrality from the Man9 glycan interaction network, and **(E)** the Endo H protection score calcualted from the time-resolved Endo H digestion experiments performed on BG505_293S. **(F)** The per-glycan normalized glycan crowding score calculated using a 44Å radius probe, which represents the peak correlation between crowding and Endo H protection in panel E.

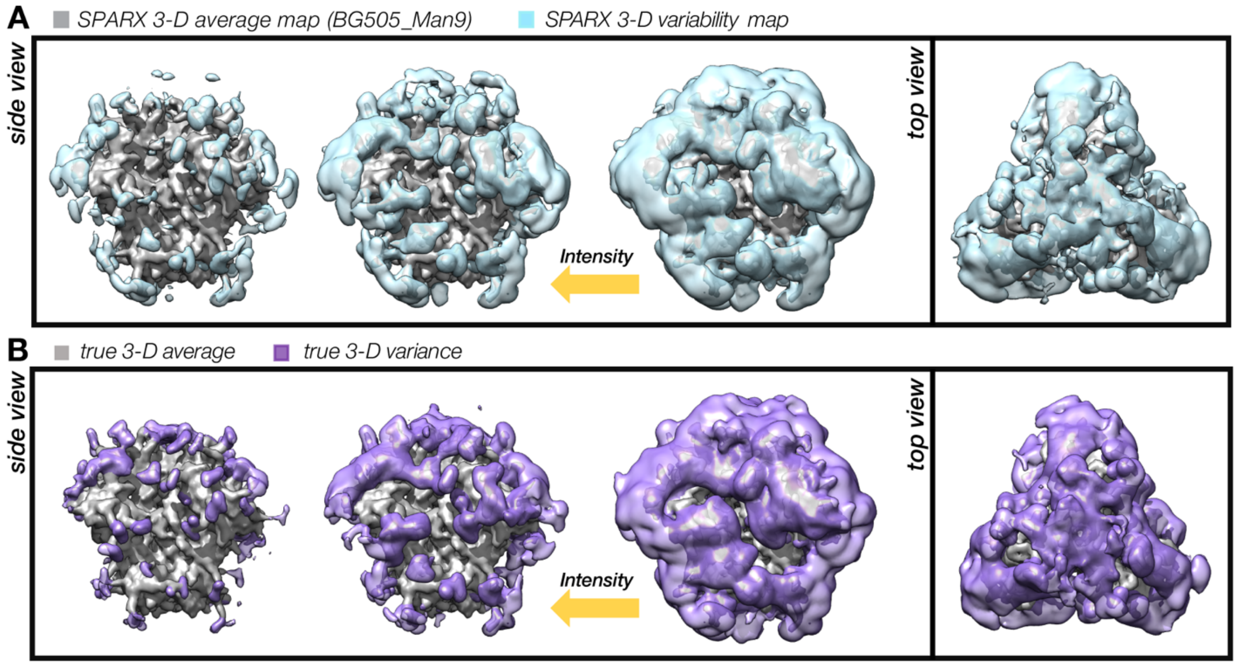


**Supplemental Figure 7 | SPARX 3-D variability analysis of simulated BG505_Man9 dataset and comparison to true 3-D variance. (A)** Side view of the Gaussian filtered (SD = 2) SPARX 3-D variability map calculated from the BG505_Man9 simulated dataset (light blue) at three intensity thresholds along with a view from the top of the trimer (R). Underneath the variability map is the SPARX 3-D variability map calculated to help diagnose the presence of potential rotational artifacts. **(B)** True per-voxel 3-D average and 3-D variance maps.

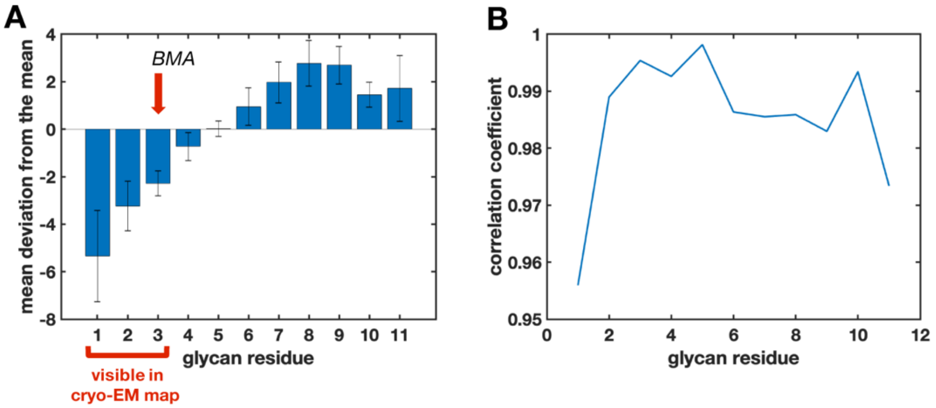


**Supplemental Figure 8 | Comparison of individual glycan residues for capturing average dynamics and relative dynamics between sites. (A)** Mean deviation from the full glycan mean rmsf for each glycan residue across all glycans in all 1000 models. **(B)** Pearson correlation coefficient between the normalized per-glycan rmsf when using the average of the entire glycan and the per-glycan rmsf when using each individual glycan residue. Both figures demonstrate that the BMA residues more closely capture the full glycan average dynamics and relative differences between sites than the first two residues. The fifth glycan residue captures the average values the closest but cannot be uniquely locate within the cryo-EM maps.

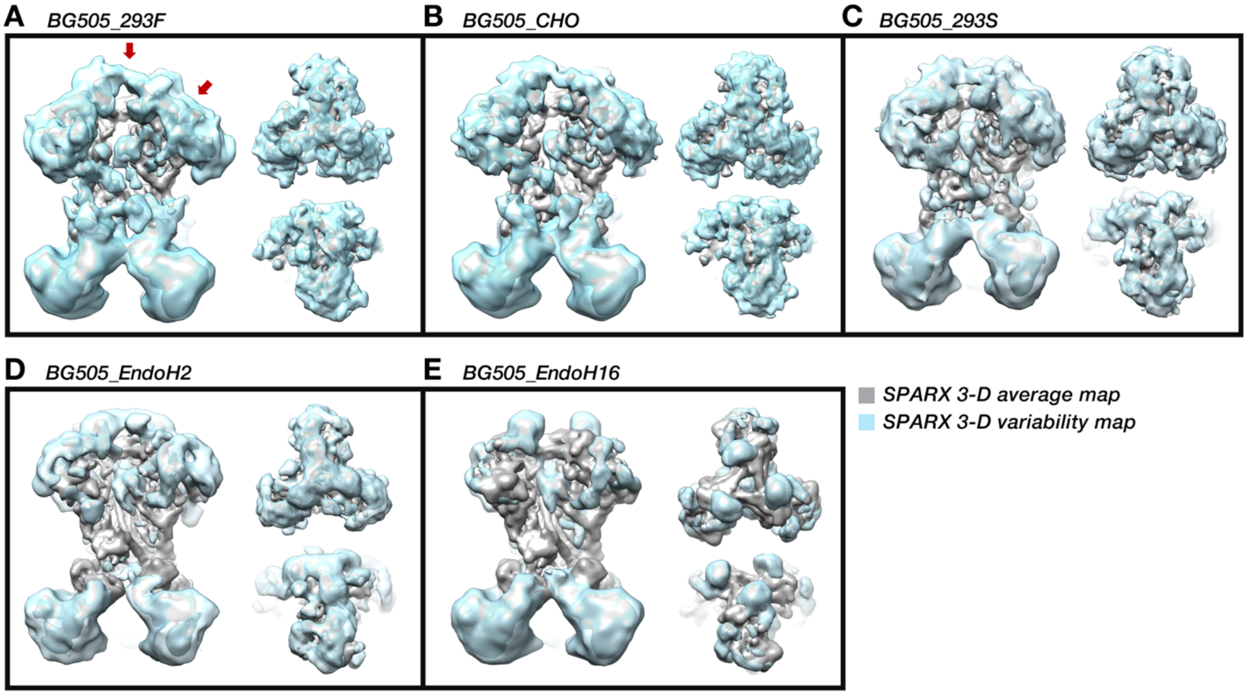


**Supplemental Figure 9 | SPARX 3-D variability analysis for all experimental cryo-EM maps.** Full side view, along with top and tilled top views of the SPARX 3-D variability maps calculated from each cryo-EM dataset presented. Red arrows indicate viewing direction. The BG505_EndoH16 showed signs of detergent aggregation in the three-fold symmetry axis of the trimer apex, which caused erroneous variance signal, so it was masked out before Gaussian filtering for clarity.

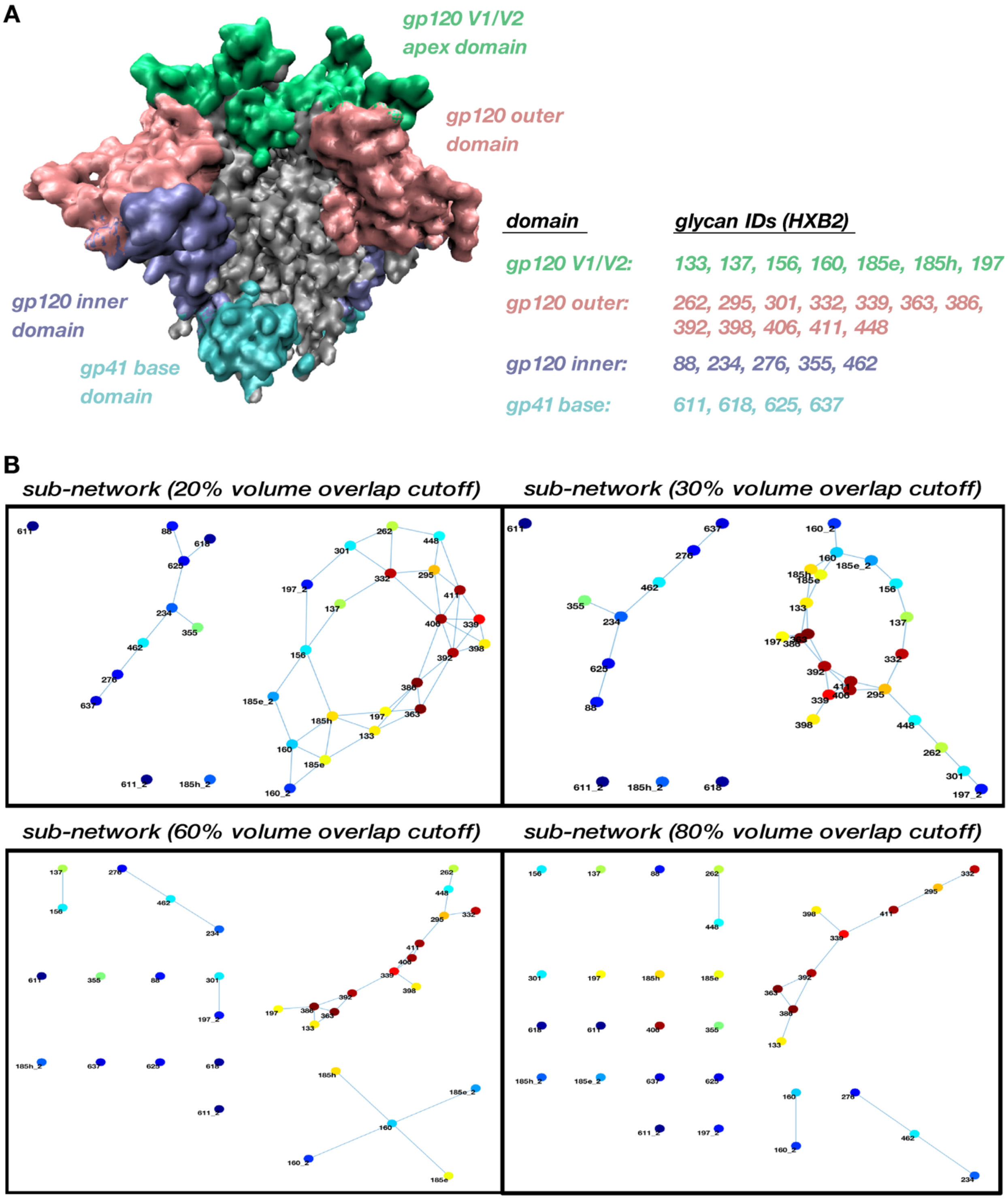


**Supplemental Figure 10 | Sub-domains in the Man9 glycan volume overlap network.** (A) Sub-domain definitions projected by constituent glycosylated ASNs on the protein surface including 10Å area surrounding them. **(B)** Sub-graphs calculated by applying a progressively stricter overlap cut-off, highlighting domain architecture. The most strongly connected glycan clusters persist as the overlap cut-off percentage increases.

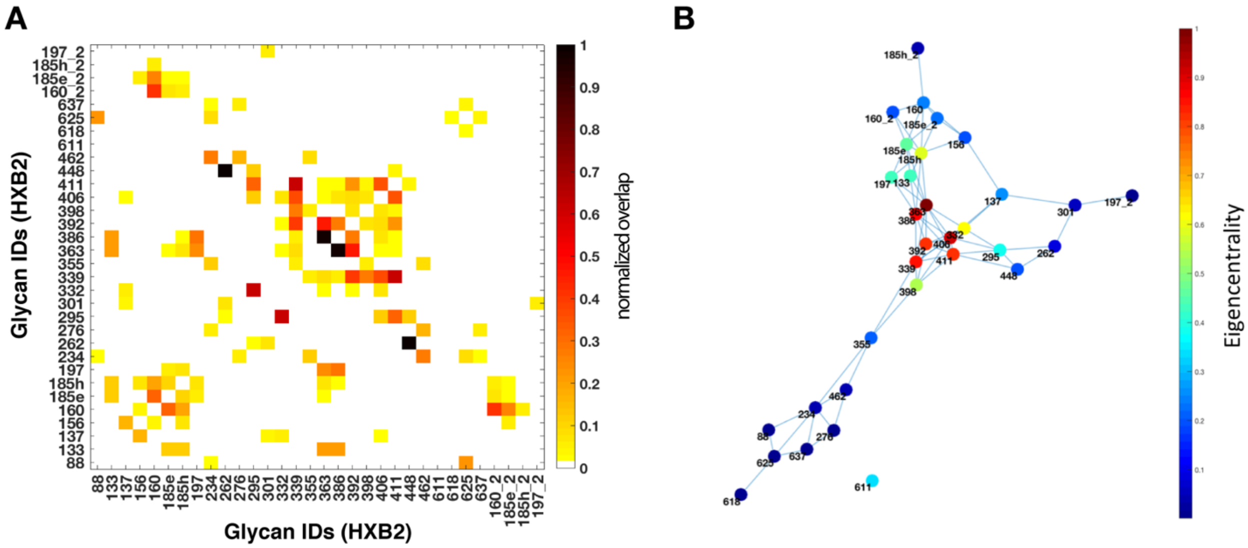


**Supplemental Figure 11 | Glycan-glycan interaction network for the Man5 HT-AM ensemble. (A)** Glycan-glycan overlap matrix calculated for the Man5 ensemble as in Figure 8. **(B)** Glycan-glycan interaction network calculated from the matrix in A. All nodes are colored by eigenvector centrality as in Figure 8.

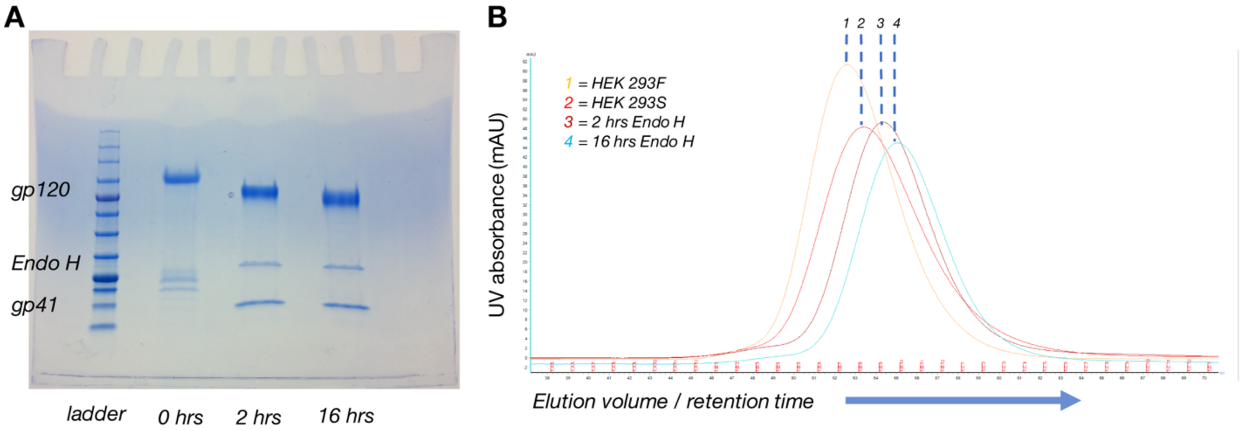


**Supplemental Figure 12| SDS PAGE gel and SEC chromatogram of Endo H digested BG505 SOSIP.664. (A)** SDS PAGE gel showing BG505 SOSIP.664 expressed and purified form HEK 293S cells after 0 hrs (L), 2hrs (M), and 16 hrs (R) of Endo H digestion. The gradual digestion of the glycan shield manifests as a gel shift in the gp120 and gp41 band. Un-digested gp41 appears as a mixed population of bands, presumably representing differentially glycosylated species. **(B)** SEC chromatogram showing a shift in retention time upon expression in HEK 293S cells (with respect to HEK 293F cells) and additional shifts due to progressive de-glycosylation by Endo H.

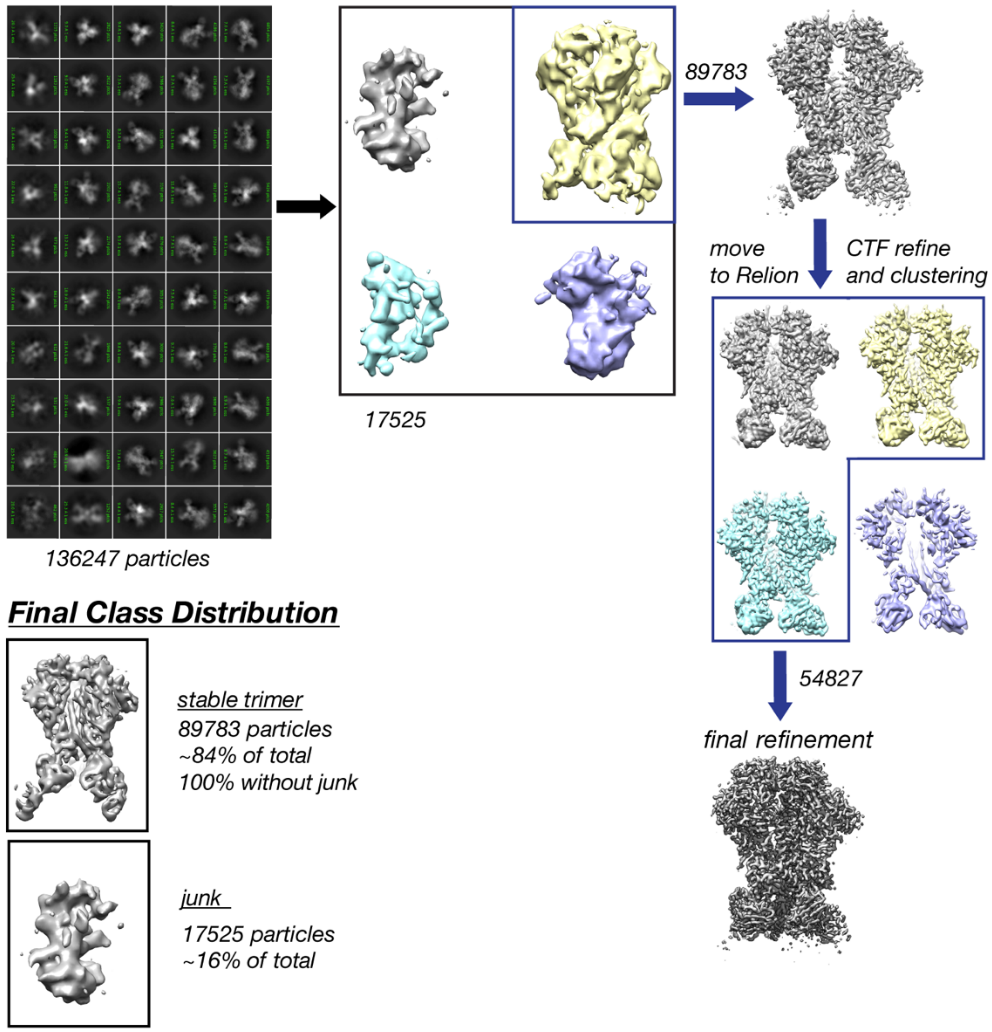


**Supplemental Figure 13 | Cryo-EM data classification tree for BG505_293F.** 2-D and initial 3-D classification and refinement in CryoSparc followed by final processing and classification in RELION. Also shown are the final class distributions among the 5 distinct classes.

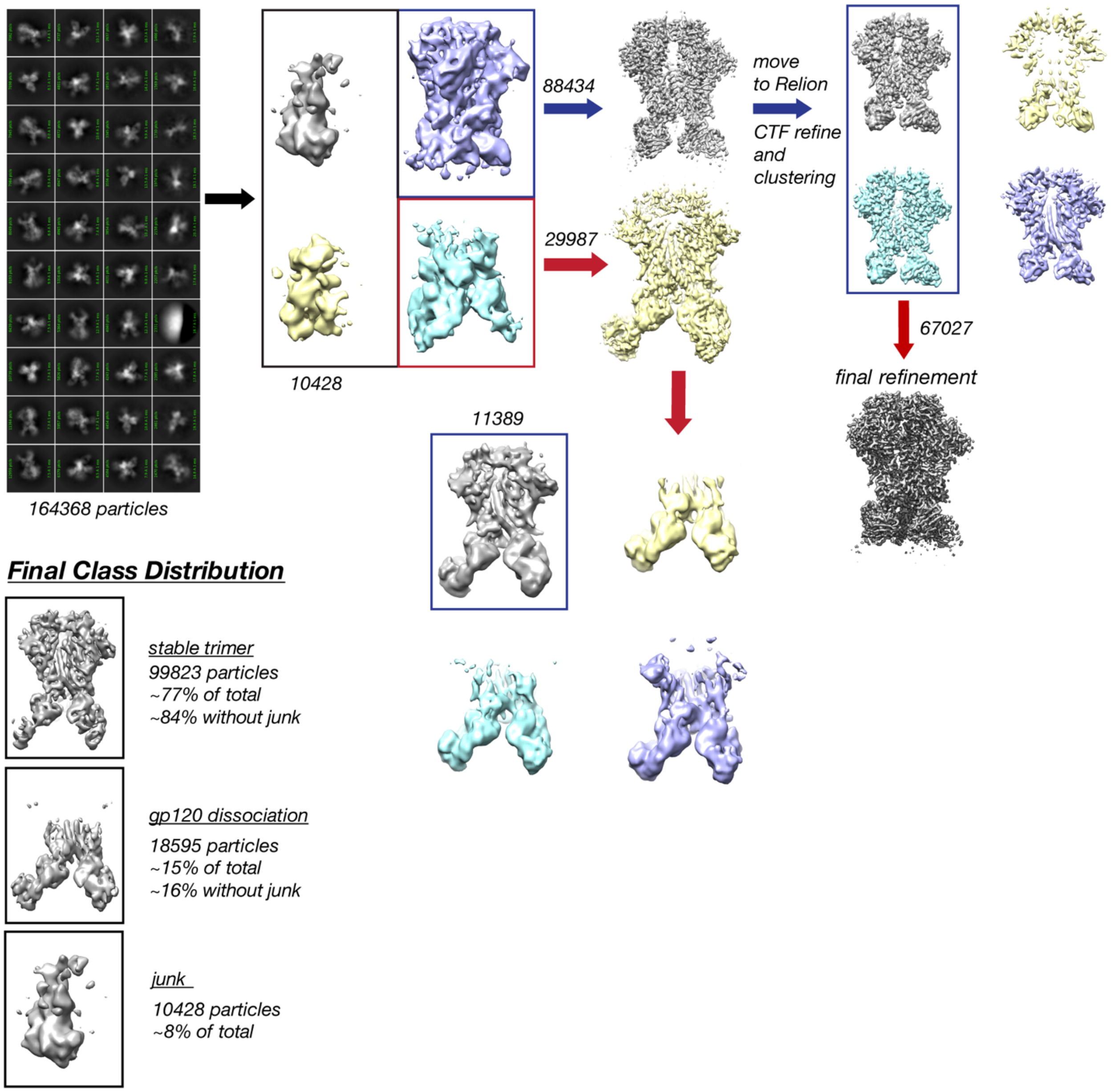


**Supplemental Figure 14 | Cryo-EM data classification tree for BG505_CHO.** 2-D and initial 3-D classification and refinement in CryoSparc followed by final processing and classification in RELION. Also shown are the final class distributions among the 5 distinct classes.

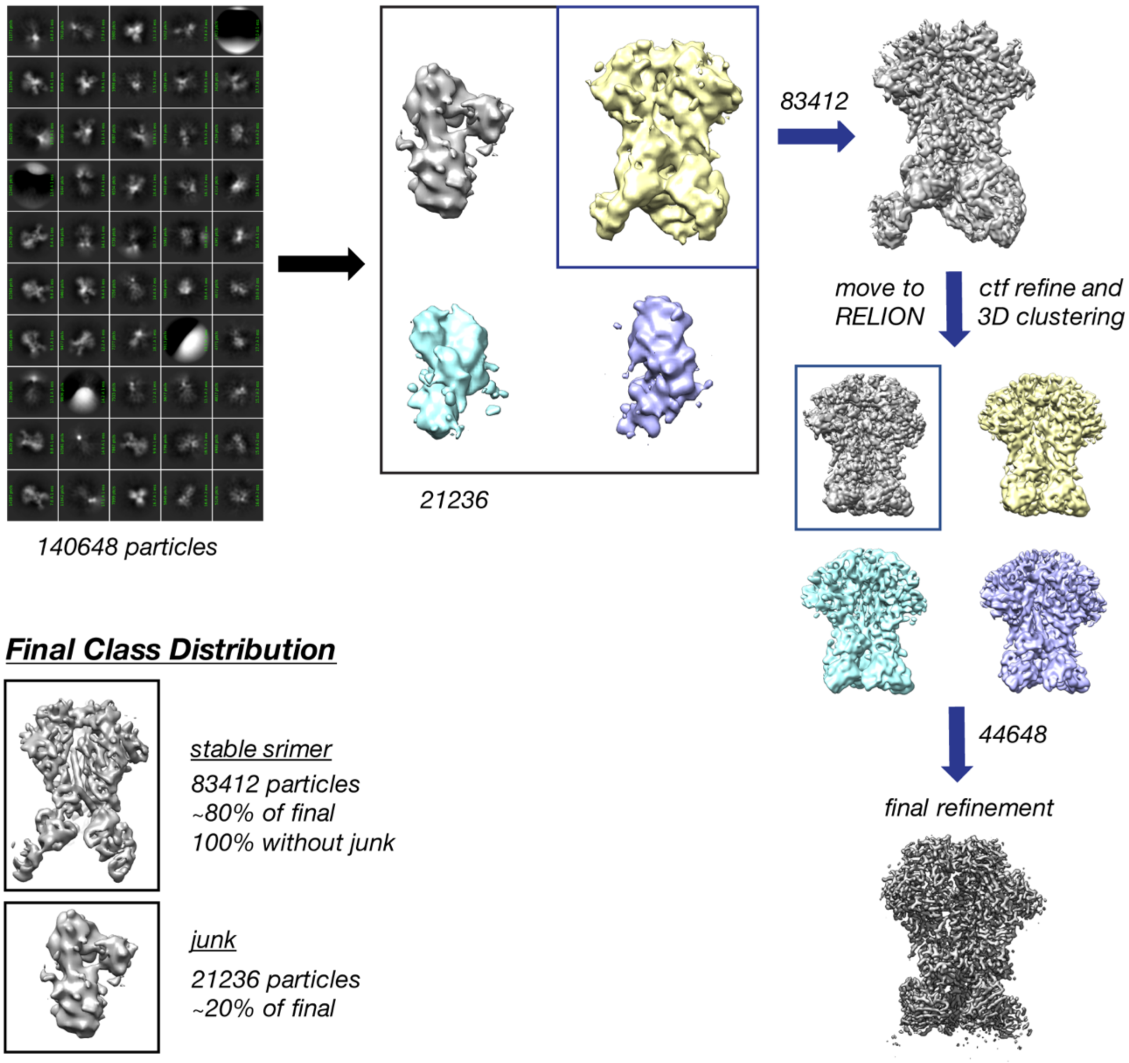


**Supplemental Figure 15| Cryo-EM data classification tree for BG505_293S.** 2-D and initial 3-D classification and refinement in CryoSparc followed by final processing and classification in RELION. Also shown are the final class distributions among the 5 distinct classes.

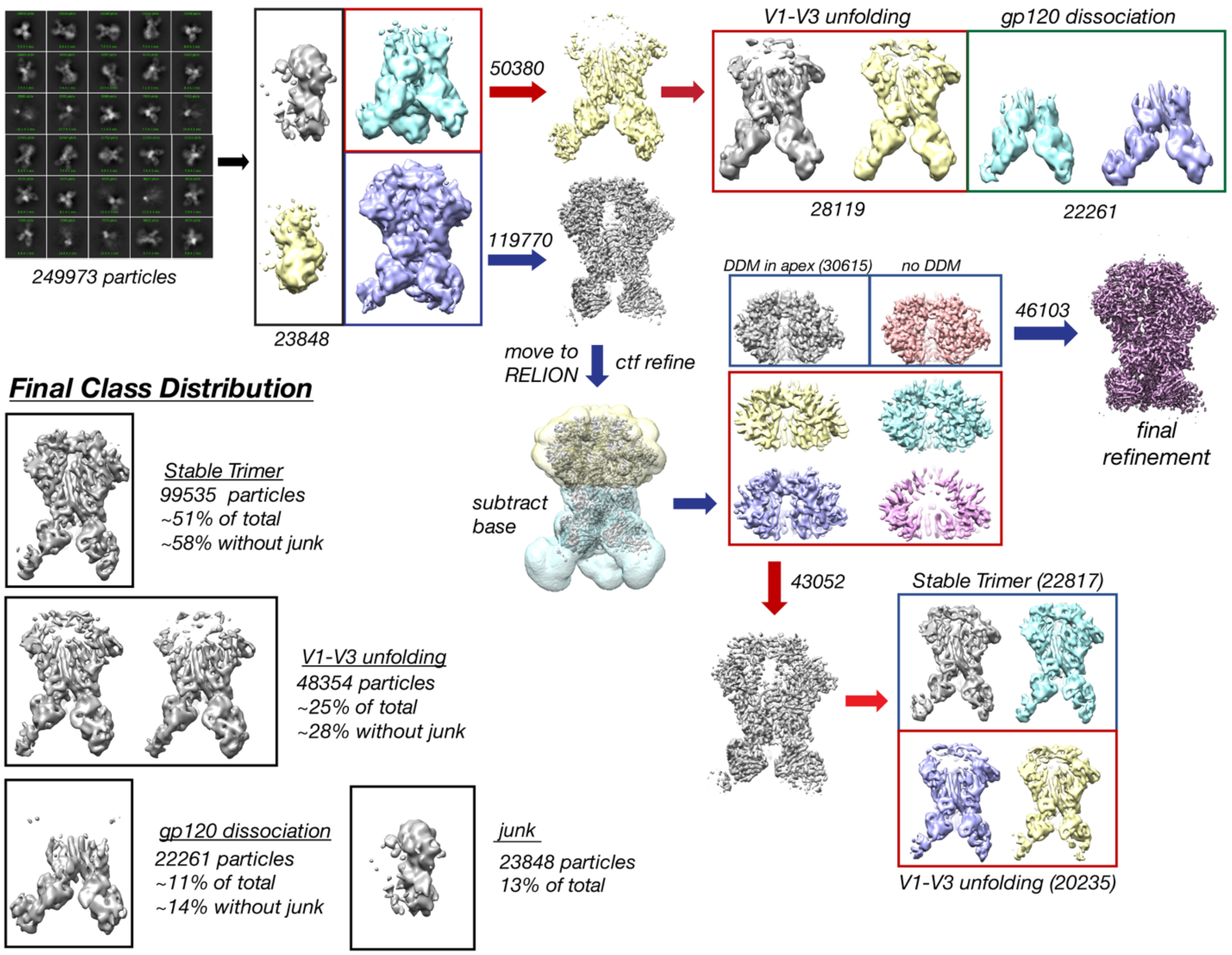


**Supplemental Figure 16 | Cryo-EM data classification tree for BG505_EndoH2.** 2-D and initial 3-D classification and refinement in CryoSparc followed by final processing and classification in RELION. Also shown are the final class distributions among the 5 distinct classes.

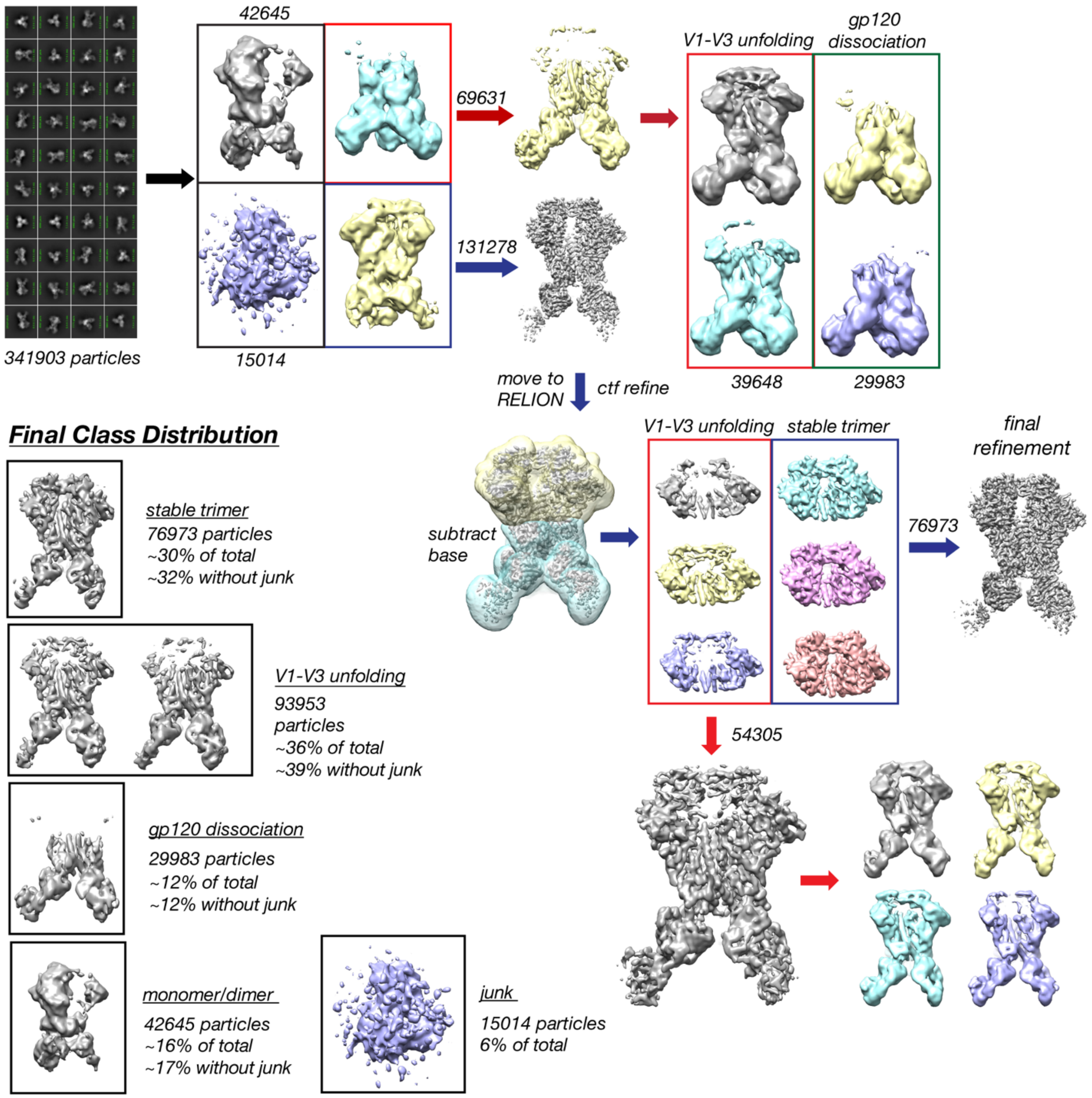


**Supplemental Figure 17 | Cryo-EM data classification tree for BG505_EndoH16.** 2-D and initial 3-D classification and refinement in CryoSparc followed by final processing and classification in RELION. Also shown are the final class distributions among the 5 distinct classes.

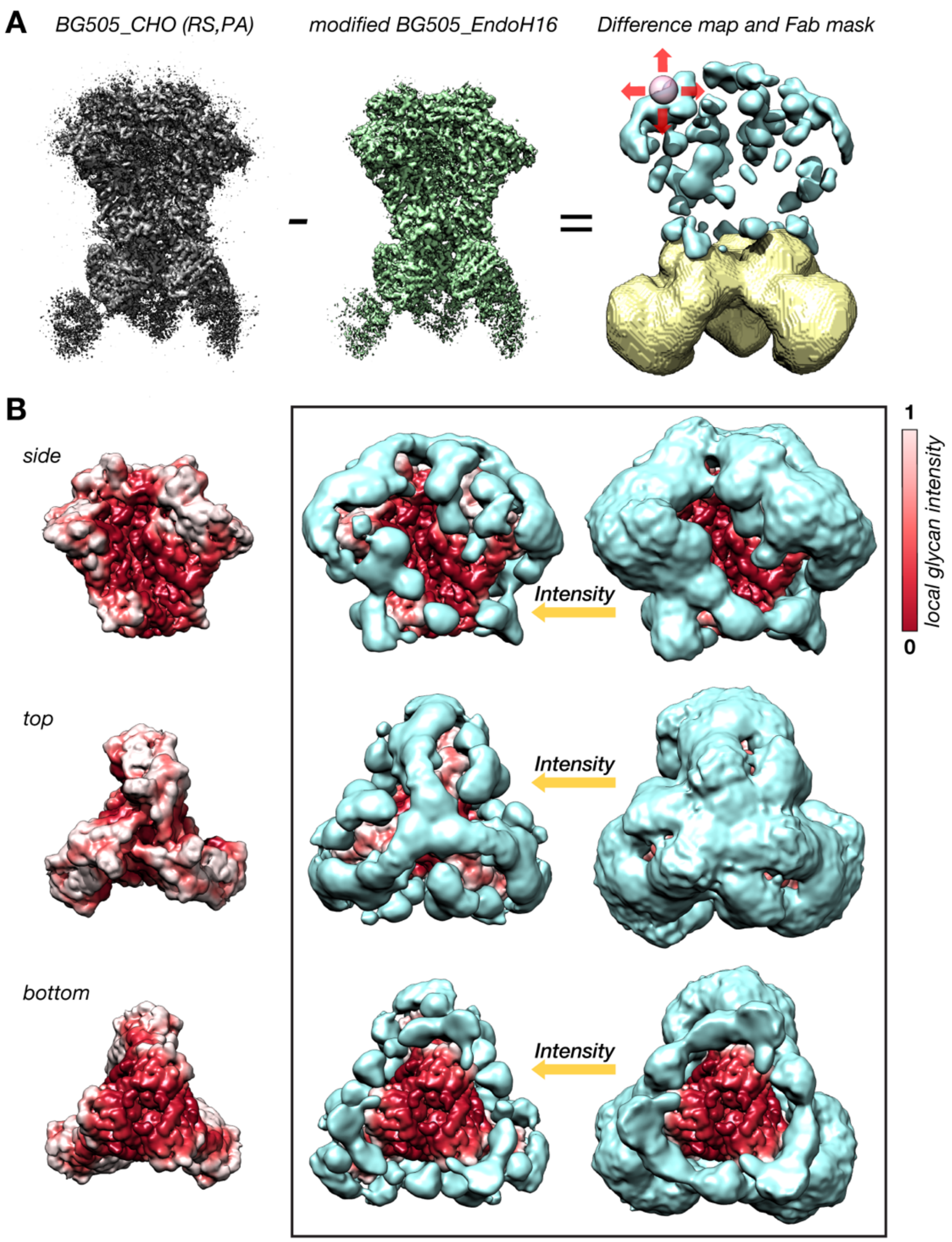


**Supplemental Figure 18 | Rolling sphere method for calculating the glycan shielding effect from cryo-EM maps.** **(A)** Schematic of the signal subtraction and masking process used to isolate the glycan shield signal. L - Unsharpened BG505_CHO map resampled (RS), and intensity equalized (PA) to the BG505_EndoH16 map. M - Unsharpened BG505_EndoH16 map with residual glycan density removed. R - Gaussian filtered difference map and the complement of the mask used to remove residual Fab signal after subtraction. Also depicted is a representation of the ~7Å radius spherical probe used to calculate local glycan intensity. **(B)** Side, top, and bottom views of the modified BG505_EndoH16 map with Fabs masked out and colored according to normalized local glycan intensity (same as in Figure 11), along with the Gaussian filtered difference map (light blue) overlaid at two intensity thresholds. The local glycan intensity was normalized such that the protein surfaces still visible at the noise-threshold (right) are red and the surfaces roughly occluded by a single glycan (left) are white.
